# Microglia from patients with multiple sclerosis display a cell-autonomous immune activation state

**DOI:** 10.1101/2024.12.10.627477

**Authors:** Tanja Hyvärinen, Johanna Lotila, Luca Giudice, Iisa Tujula, Marjo Nylund, Sohvi Ohtonen, Flavia Scoyni, Henna Jäntti, Sara Pihlava, Heli Skottman, Susanna Narkilahti, Laura Airas, Tarja Malm, Sanna Hagman

## Abstract

Aberrant and sustained activation of microglia is implicated in the progression and severity of multiple sclerosis (MS). However, whether intrinsic alterations in microglial function impact the pathogenesis of this disease remains unclear.

We conducted transcriptomic and functional analyses of microglia-like cells (iMGLs) differentiated from induced pluripotent stem cells (iPSCs) from patients with MS (pwMS) to answer this question.

We generated iPSCs from six pwMS showing increased microglial activity via translocator protein (TSPO)-PET imaging. We demonstrated that the differentiated iMGL transcriptional profile resembled the microglial signature found in MS lesions. Importantly, compared with healthy controls, MS iMGLs presented cell-autonomous differences in their regulation of inflammation, both in the basal state and following inflammatory lipopolysaccharide challenge. Through transcriptomic profiling, we showed that MS iMGLs display increased expression of genes known to be upregulated in MS microglia. Furthermore, upregulated genes in MS iMGLs were associated with immune receptor activation, antigen presentation, and the complement system, with known MS implications. Finally, functional analyses indicated that the transcriptional changes in MS iMGLs corresponded with alterations in the secretion of inflammatory cytokines and chemokines and increased phagocytosis.

Together, our results provide evidence of putative cell-autonomous microglial activation in pwMS and identify transcriptomic and functional changes that recapitulate the phenotypes observed *in vivo* in microglia from pwMS. These findings indicate that MS disease-specific iPSCs are valuable tools for studying disease-specific microglial activation *in vitro* and highlight microglia as potential therapeutic targets in MS.

## Introduction

Multiple sclerosis (MS) is a chronic autoimmune disease of the central nervous system (CNS) characterized by focal demyelinated lesions, inflammation and neurodegeneration^1^. Both environmental and lifestyle factors along with genetic predisposition influence the risk of developing MS^2^. Recent genome-wide association studies (GWASs) have identified more than 200 susceptibility variants for MS^3^. Interestingly, these identified susceptibility variants are enriched not only in peripheral immune cells but also in microglia, which are the brain-resident immune cells, suggesting a role in disease onset^3^. Furthermore, newly discovered severity variants link for the first time, faster MS progression and increased cortical pathology to CNS cell functions. This finding underscores the importance of CNS-specific pathological mechanisms in modulating the severity of MS^4^.

From the onset of the disease, underlying pathological processes within the CNS contribute to the development of disability^5–7^. Disease progression is associated with the presence of chronic active lesions characterized by a hypocellular centre surrounded by activated microglia at the lesion edge^1,8,9^. Many of these lesions can also be identified by the presence of iron-laden microglia and macrophages using brain magnetic resonance imaging (MRI) which can predict more severe tissue damage and worse clinical outcomes^10,11^. Similarly, positron emission tomography (PET) imaging of the mitochondrial 18-kDa translocator protein (TSPO) ligand has been utilized to study microglial involvement *in vivo*^12^. PET studies have shown that microglial activity precedes signs of neurodegeneration^13^ and is associated with faster disease progression^14^. Thus, this evidence suggests that microglia play a critical role in the immunopathological processes of MS. However, the underlying mechanisms and their contributions to the chronic stages remain uncertain.

Microglia are highly dynamic, phagocytic cells that constantly survey and maintain tissue homeostasis in the CNS. Under pathological conditions, they react and transform into different functional states according to context-dependent signals, including the local immune environment and the functions of other cells^15–17^. However, over time, the initial resolving immune response can escalate into a vicious cycle of chronic neuroinflammation with a potential loss of homeostatic functions and increased production of neurotoxic factors, including proinflammatory cytokines, chemokines, nitric oxide and reactive oxygen species^18^. Aberrant and sustained inflammatory responses of microglia have been documented in experimental animal models of MS and histopathological postmortem brain tissues from patients with MS (pwMS)^19–23^. Studies have reported the loss of homeostatic microglial markers and the presence of activated microglia in various MS lesion types and normal-appearing white matter (NAWM) from autopsy tissues of pwMS ^24,25^. This microglial activation becomes more pronounced with increasing disease duration^24^. Notably, clusters of activated microglia, known as microglial nodules, are present in the NAWM before lesion formation^25,26^.

Single-cell (sc) and single-nucleus (sn) RNA-sequencing (RNA-seq) studies of postmortem brain tissue from pwMS have provided valuable insights into disease-associated microglial states^20,21,25,27–29^. These studies identified gene signatures involved in immune regulation, lipid metabolism, iron homeostasis, phagocytosis, the complement system and antigen presentation^20,21,25,27^. A recent seminal study described two distinct phenotypes of microglia inflamed in MS (MIMS), one referred to as MIMS-foamy, which is involved in the phagocytosis and clearance of myelin, and the other, MIMS-iron, which is characterized by the enrichment of iron-and complement-related genes^20^. Despite these advancements, it remains unknown whether MS microglia exhibit cell-autonomous alterations that contribute to disease susceptibility and disease processes, such as lesion formation or progression. Understanding microglial dysfunction in MS is crucial for developing targeted therapies.

Stem cell technology represents a complementary research tool to traditional disease models. Human induced pluripotent stem cells (iPSCs) can be generated from pwMS, providing unique opportunities to study this disease^30^. To date, studies of MS iPSCs have revealed several key findings: the senescence of neural progenitors^31^, deficits in oligodendrocyte progenitor cell migration^32^ and differentiation in an inflammatory milieu^33^, increased inflammatory activation of astrocytes^34^, altered metabolic function of astrocytes^35^, and disrupted barrier function of microvascular endothelial cells^36^. In recent years, pioneering studies have reported methods to produce microglia-like cells (iMGLs) from iPSCs, addressing the previously unmet need for *in vitro* models of microglial phenotypes in neurological disorders, including MS^37–40^. In this study, we aimed to determine whether MS patient-derived iMGLs exhibit cell-intrinsic alterations in their homeostatic and immune activation states. We investigated these properties by generating iPSC lines from six pwMS showing microglial activation using TSPO-PET imaging, differentiated them into iMGLs and compared them to healthy controls (HCs) by performing RNA-seq and functional assays, including secretome and phagocytosis analyses. We found that MS patient-derived iMGLs displayed a distinct inflammatory phenotype at the transcriptome level, which translated into functional changes. These changes included alterations in cytokine and chemokine secretion and increased phagocytic activity. These findings suggest that patient-specific iMGLs represent an exceptional tool for modelling microglial dysfunction in MS and may inform therapeutic strategies that focus on targeting microglia.

## Materials and methods

The detailed methods are available in the Supplementary material.

### Subjects and procedures

Six pwMS were recruited during 2020 from the Turku MS PET cohort (Table 1). Two patients had relapsing-remitting MS (RRMS), and four had secondary progressive MS (SPMS). The Turku MS PET cohort consists of over 150 patients with different stages of MS who have participated in different MS PET studies at the Turku PET centre and have been examined using PET, MRI and clinical evaluation between 2009 and 2024. All participants were interviewed and clinically examined by a neurologist. The Expanded Disability Status Scale (EDSS) score^41^ was assessed for patients. The inclusion criteria were a confirmed MS diagnosis according to the 2010 McDonald criteria^42^, a previous PET scan using the [^11^C]PK11195 radioligand and female sex. Patients provided informed consent, after which a blood sample was obtained for iPSC reprogramming. For comparison, six similarly imaged age-and sex-matched healthy controls were included for MRI and PET imaging analyses (Table 1). The study protocol was approved by the Ethics Committee of the Hospital District of Southwest Finland (Dnro: 48/1801/2019) and the study was conducted in accordance with the principles of the Declaration of Helsinki.

**Table 1.** Demographic, clinical and radiological characteristics of study subjects.

| Variable | HC | MS | MS vs HC *p* value |
| --- | --- | --- | --- |
| n | 6 | 6 |  |
| Sex, n |  |  |  |
| Females | 6 | 6 |  |
| Age | 50.0 (44.7–60.1) | 48.6 (45.7–56.7) |  |
| Disease type |  |  |  |
| RRMS |  | 2 |  |
| SPMS |  | 4 |  |
| Duration, years |  | 16.9 (12.9–28.4) |  |
| EDSS |  | 3.5 (2.5–7.5) |  |
| MSSS |  | 3.7 (1.2–8.1) |  |
| ARR |  | 0.3 (0.2–0.8) |  |
| Treatment |  |  |  |
| Untreated |  | 3 |  |
| Interferon-β | | 1 | |
| Dimethylfumarate |  | 1 |  |
| Glatiramer acetate |  | 1 |  |
| MRI measures (volumes, cm<sup>3</sup>) |  |  |  |
| T1 |  | 10.5 (0.1–28.1) |  |
| T2 |  | 16.6 (0.3–43.3) |  |
| Whole-brain (PF) | 0.868 (0.836–0.916) | 0.803 (0.769–0.879) | **0.041** |
| NAWM (PF) | 0.354 (0.325–0.439) | 0.309 (0.281–0.381) | 0.132 |
| GM cortex (PF) | 0.333 (0.308–0.348) | 0.305 (0.281–0.328) | 0.065 |
| Thalamus (PF) | 0.012 (0.011–0.015) | 0.009 (0.008–0.012) | **0.013** |

Data are expressed as median (min–max) unless specified otherwise. Abbreviations: HC, healthy control; MS, multiple sclerosis; RRMS, relapsing-remitting multiple sclerosis; SPMS, secondary progressive multiple sclerosis; EDSS, Expanded Disease Status Scale; MSSS, Multiple Sclerosis Status Scale; ARR, annualized relapse rate; PF, parenchymal fraction; NAWM, normal-appearing white matter; GM, grey matter. Mann-Whitney U-test was used to compare MS vs HC. Here, $p < 0.05$ is considered significant and values are bolded.

### Generation and maintenance of iPSCs

The control iPSC lines UTA.04511.WT^43^, UTA.04602.WT^43^, UTA.10902.EURCCs^44^ and UTA.11311.EURCCs^45^ were previously generated and characterized at Tampere University with the approval of the Ethics Committee of Wellbeing Services County of Pirkanmaa (R08070, R12123). For this study, supportive ethical statements for producing MS patient-derived iPSC lines (48/1801/2019) and for culturing and differentiating iPSCs for neuronal research (R20159) were obtained from the Ethics Committee of the Hospital District of Southwest Finland and the Wellbeing Services County of Pirkanmaa, respectively. Informed consent was obtained from all the subjects. The iPSC lines are listed in Supplementary Table 1. Six MS iPSC lines were generated as previously described^46^. One of the MS iPSC lines can also be found at the Human Pluripotent Stem Cell Registry (https://hpscreg.eu) with the hPSCreg name TAUi008-A. IPSCs were produced from peripheral blood mononuclear cells (PBMCs) using the CytoTune™-iPS 2.0 Sendai Reprogramming Kit (Thermo Fisher Scientific). The iPSC lines were derived on a feeder layer of mitomycin C-inactivated human foreskin fibroblasts (CRL-2429™, ATCC). All iPSC lines were subsequently maintained as previously described^44^ in feeder-free cultures on 0.6 µg/cm^2^ recombinant human laminin-521 (LN521, Biolamina)-coated cell culture plates in Essential 8™ Flex medium (E8 flex, Thermo Fisher Scientific). IPSCs were passaged enzymatically with TrypLE™ Select Enzyme and Defined Trypsin Inhibitor (both from Thermo Fisher Scientific) in the presence of 10 µM ROCK inhibitor (ROCKi, Y-27632, StemCell Technologies) twice a week.

### Differentiation of iMGLs

MS (n=6) and HC (n=4) iPSCs were differentiated into iMGLs according to the protocol reported by Konttinen et al.^37^, with light modifications. On Day 0, iPSCs were seeded at a density of 6,200–29,000 cells/cm^2^ on Matrigel (Corning)-coated dishes and cultured under low-oxygen conditions (5% O_2_, 5% CO_2_, 37 °C) until Day 4. On Days 0 and 1, the cells were maintained in E8 flex -medium containing 5 ng/ml BMP4, 25 ng/ml activin A (both from Peprotech) and 1 µM CHIR 99021 (Axon). The medium contained ROCKi for the first two days of culture (Day 0: 10 µM and Day 1: 1 µM). From Day 2 until Day 8, the cells were cultured in basal medium containing DMEM/F-12 without glutamine, 1X GlutaMAX, 543 mg/l sodium bicarbonate (all from Thermo Fischer Scientific), 14 µg/l sodium selenite, 64 mg/l L-ascorbic acid (both from Sigma–Aldrich) and 0.5% penicillin/streptomycin (P/S). On Days 2 and 3, the basal medium was supplemented with 100 ng/ml FGF2, 50 ng/ml VEGF (both from Peprotech), 10 µM SB431542 and 5 µg/ml insulin (both from Sigma–Aldrich). From Day 4 onwards, the cells were maintained in a normoxic incubator. From Day 4 until Day 8, the basal medium was supplemented with 50 ng/ml FGF2, 50 ng/ml VEGF, 50 ng/ml TPO, 50 ng/ml IL-6, 10 ng/ml SCF, 10 ng/ml IL-3 (all from Peprotech) and 5 µg/ml insulin (Sigma-Aldrich) and changed daily. On Day 8, floating erythromyeloid progenitor cells (EMPs) were harvested and plated at a density of 64,000 cells/cm^2^ in ultralow attachment dishes (Corning). Cultures were maintained in basal medium containing Iscove′s modified Dulbecco′s medium (IMDM, Thermo Fischer Scientific), 10% heat-inactivated fetal bovine serum (FBS, Sigma– Aldrich) and 0.5% P/S supplemented with 5 ng/ml MCSF, 100 ng/ml IL-34 (both from Peprotech) and 5 µg/ml insulin. Beginning on Day 10, the medium supplemented with 10 ng/ml MCSF and 10 ng/ml IL-34 was changed every other day until the final plating on Day 16, when the cells were plated for experiments, after which 50% of the medium was changed daily. The experiments included immunocytochemistry, RNA-seq, RT–qPCR, western blot, secretome profiling and phagocytosis assays, and they were performed on Days 21-23 in the basal state or after inflammatory treatments.

### RNA sequencing and analysis

The iMGLs were cultured on 6-well tissue-culture treated plates at a density of 400,000 cells/well and stimulated with LPS for 24 h. RNA was extracted from vehicle and LPS-treated iMGLs with a NucleoSpin RNA Kit (Macherey-Nagel) according to the manufacturer’s protocol, and the samples from two wells were pooled. RNA integrity was analysed with a DNF-471 RNA Kit (15 nt) and Fragment Analyzer (both from Agilent) according to the manufacturer’s instructions. Libraries were prepared with a CORALL Total RNA-Seq V2 Library Prep Kit with RiboCop (Lexogen). The quality of the prepared cDNA libraries was analysed with a High Sensitivity DNA Analysis Kit (Agilent). Sequencing was performed using an Illumina NextSeq500 sequencer with an Illumina NextSeq 500/550 High Output Kit v2.5, 75 cycles -kit (both from Illumina).

### Analysis of RNA sequencing data

The bulk RNA sequencing dataset was analysed using a series of established bioinformatic techniques. The raw count data were obtained with the nf-core rnaseq pipeline^47^. The raw count data were initially prepared by removing low-quality reads and normalizing by library size using the trimmed mean of M-values (TMM) method implemented in the edgeR package^48^. Quality control was performed by employing the similarity measure implemented in the StellarPath package^49^, and the result was visualized through multidimensional scaling (MDS) to identify potential batch effects and outliers. This analysis revealed that the cell line was a significant confounding factor influencing the samples’ expression profiles. Therefore, cell line information was included as a covariate in the downstream differential expression analysis. Gene set variation analysis (GSVA) was performed using single-cell-specific markers derived from PanglaoDB^50^ to identify the cell types contributing to each sample. The ssGSEA method was chosen for GSVA to calculate enrichment scores representing the degree of activation of each cell type-specific gene set in each sample. The same operation was also performed to compare our samples with the samples described by Abud et al.^38^. Subsequently, we also performed this task with the markers reported by Absinta et al.^20^ to determine the type of microglia present in our samples. The differential expression analysis was then conducted using the limma package^51^ with a contrast matrix designed to compare specific conditions of interest. This analysis aimed to identify genes whose expression levels were significantly altered between the compared conditions. Finally, a pathway analysis was performed using clusterProfiler^52^ and msigdbr^53^ to identify enriched pathways and biological processes associated with the observed changes in gene expression. This step provided insights into the biological functions and pathways affected by the experimental conditions.

## Results

### Brain TSPO-PET imaging reveals microglial activation in pwMS

We recruited six pwMS, including four patients with SPMS and two with RRMS disease. The clinical and radiological characteristics of the subjects are summarized in Table 1. The median disease duration was 16.9 (min–max 12.9–28.4) years representing generally a more advanced disease state. At the time of imaging, the median Expanded Disability Status Scale (EDSS) score was 3.5 (range 2.5–7.5). Half of the patients were untreated at the time of imaging or sampling. The patients were imaged with conventional MRI and PET (Fig. 1A), and the results were compared with those of age-and sex-matched healthy control subjects. MRI scans of the pwMS showed ongoing tissue damage and inflammation, with median (range) T1 and T2-detected lesion loads of 10.5 (0.1–28.1) and 16.6 (0.3–43.3), respectively (Table 1). Compared with HCs, brain volumetric measures indicated that pwMS presented with brain atrophy, as evidenced by significantly lower whole-brain and thalamic volumes but no changes in NAWM or cortical grey matter (GM) volumes (Table 1). PET imaging with the TSPO-binding radioligand [^11^C]PK11195^54^ was performed to evaluate ongoing microglial activation. The distribution volume ratio (DVR), which is used to quantify TSPO binding, indicated significantly greater innate immune cell activation within the whole brain and NAWM of the pwMS than in HCs (1.222 vs. 1.169, median, *p* = 0.026 and 1.241 vs. 1.148, *p*=0.0043), but no changes were observed in the thalamic region (Fig. 1B). These findings highlight ongoing microglial activation and its potential role in the pathology of MS.

**Figure 1.**
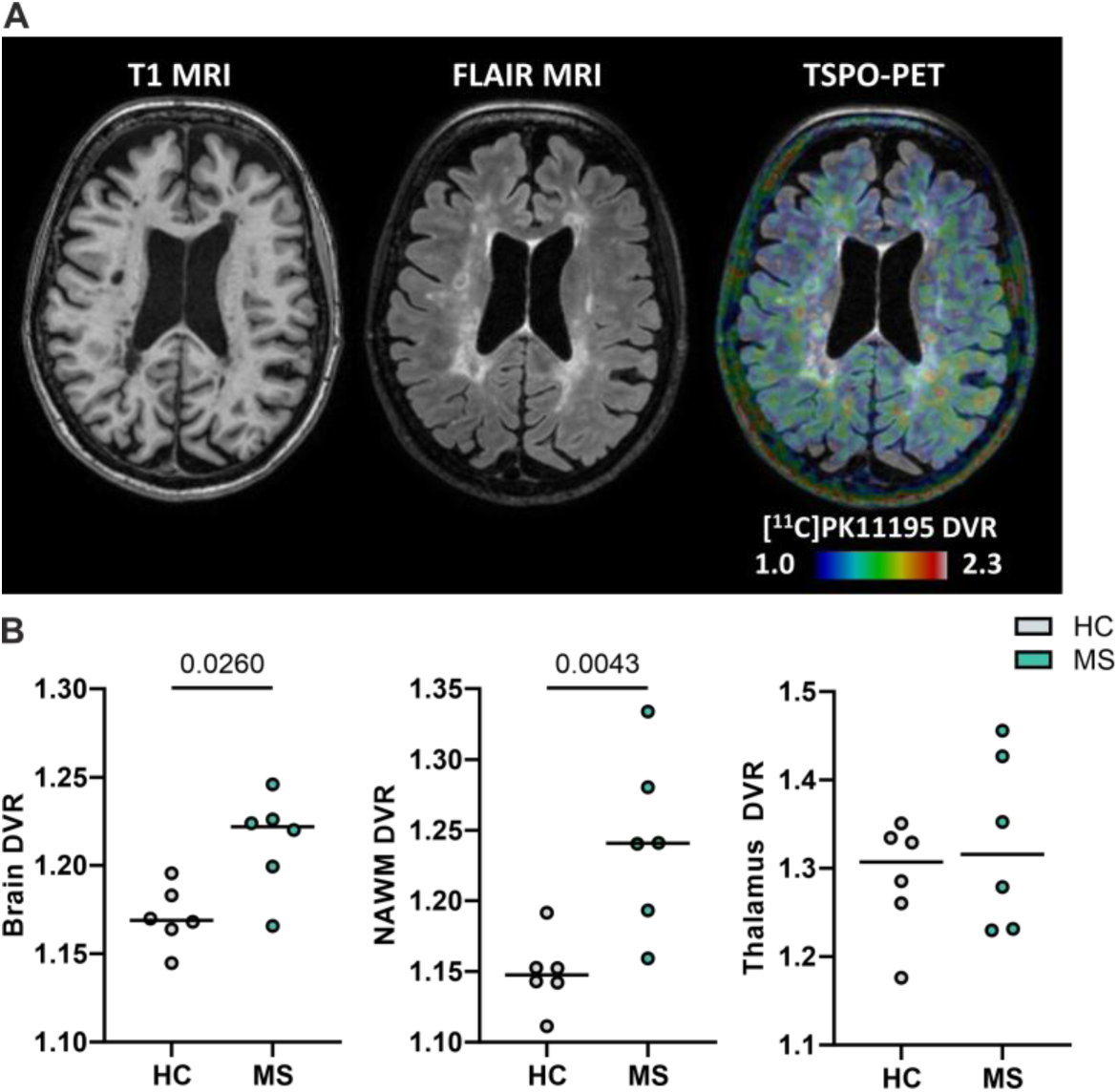
^11^C-PK11195-PET brain imaging of pwMS and HCs. (**A**) T1-weighted, FLAIR and TSPO-PET images of a patient with MS show a significant lesion load and increased microglial activation, as evidenced by increased [^11^C]PK11195 binding in the NAWM and perilesional areas. The colour bar shows the dynamic range of the DVR in the PET image. (**B**) PET-measured brain DVR, NAWM DVR and thalamus DVR of the HCs (*n = 6*) and pwMS (*n* = 6) included in this study. The data are presented as single datapoints and medians. Significance was determined with the Mann–Whitney U test.

### Differentiated iMGLs exhibit microglial features and mimic the transcriptional phenotypes observed in MS lesions

To investigate the MS-specific microglial phenotype and elucidate the potential disease mechanisms of microglial activation, we generated iPSC lines from these six pwMS. The iPSCs were successfully generated from PBMCs using Sendai virus reprogramming, and newly established iPSC lines were characterized based on their expression of pluripotency markers and trilineage differentiation capacity (Supplementary Fig. 1). A quality control assessment confirmed that the iPSC lines were free of viral transgenes and mycoplasma (Supplementary Fig. 2). The iPSCs presented a normal karyotype and correct cell line identity, as verified via short tandem repeat (STR) analysis (Supplementary Fig. 2). For comparison, four iPSC lines derived from healthy controls were also utilized (Supplementary Table 1).

For the differentiation of iMGLs we employed a previously published protocol by Konttinen and colleagues^37^ that generates microglia-like cells by mimicking *bona fide* microglia originating from the yolk sac (Fig. 2A). All six MS and four HC iPSC lines differentiated into iMGLs and presented a typical ramified morphology (Fig. 2B; Supplementary Fig. 3A). Immunofluorescence staining revealed a high percentage of ionized calcium-binding adapter molecule 1 (Iba1) expression in iMGL, with median values ranging from 72% to 90% across the lines, and showed ubiquitous expression of the microglia-specific proteins transmembrane protein 119 (TMEM119) and the purinergic receptor P2RY12 (Fig. 2C and D, and Supplementary Fig. 3B and C). The microglial identity of differentiated cells was further confirmed via an RNA-seq analysis of all four HC and six MS iPSC lines, including technical replicates for two of the HC lines. We challenged the iMGLs by exposing them to lipopolysaccharide (LPS), a common inflammatory stimulant used to activate microglia, and included LPS-stimulated iMGLs in our dataset. Multidimensional scaling analysis of gene expression profiles showed a clear separation between vehicle-and LPS-treated iMGLs (Fig. 2E). A comparative analysis with the publicly available dataset from Abud et al.^38^ showed that our iMGLs closely resembled the microglial cells in their dataset while diverging from CD14^+^ and CD16^+^ monocytes (Fig. 2E). This finding was further validated by a cell type signature analysis, which showed that the expression profiles of our iMGLs were enriched for markers representative of microglia (Fig. 2F).

**Figure 2.**
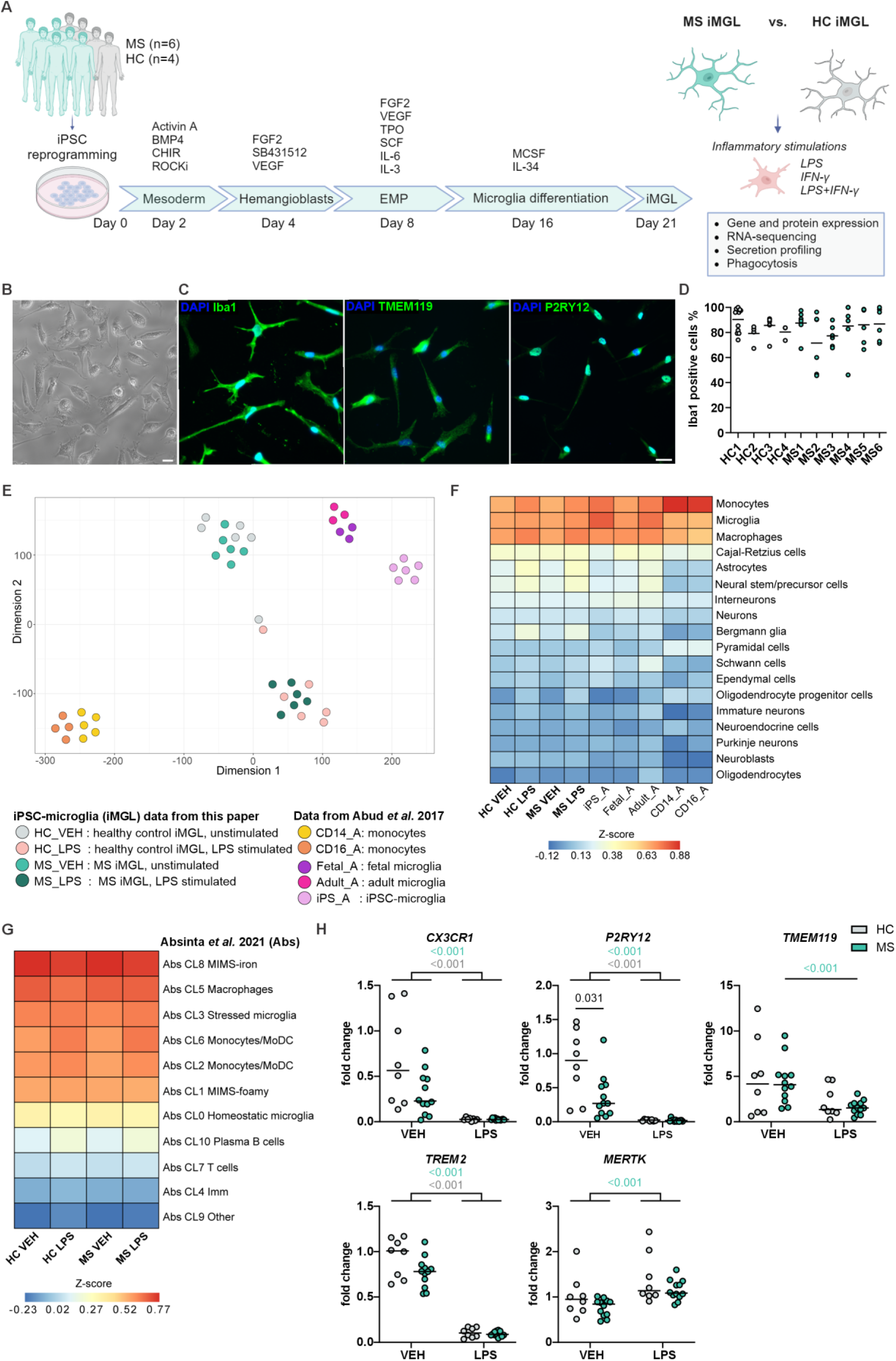
Generation and characterization of HC and MS iPSC-derived iMGLs. (**A**) Schematic overview of iMGL differentiation through erythromyeloid progenitors (EMP) and the experimental setup performed on HC and MS iMGLs. Representative images of (**B**) live cells and (**C**) immunofluorescence staining for Iba1, TMEM119 and P2RY12. Scale bar = 20 µm. (**D**) Percentages of Iba1-positive cells in immunofluorescence images, *n* = 3–15 images, with 1–4 independent differentiations per cell line. The data are presented as single datapoints and medians. No significant differences were observed between the cell lines using the Kruskal–Wallis test with Dunn’s post hoc test. (**E**) T-distributed stochastic neighbor embedding (t-SNE) plot showing the similarity between the gene expression profiles of our vehicle-and LPS-stimulated HC and MS iMGLs, and iPSC-, fetal-and adult-derived microglia and monocytes from the Abud et al.^38^ dataset. (**F**) Heatmap depicting the results of the cell type signature analysis of our iMGLs (data were merged from 6 MS and 6 HC samples, including technical replicates for two of the HC lines) and microglia and monocytes from the Abud et al.^38^ dataset, with the cell signature obtained from PanglaoDB^50^. (**G**) Heatmap depicting the cell type signature analysis of our iMGLs (data were merged from 6 MS and 6 HC samples), with the cell signature from the single-cell RNA sequencing clusters reported by Absinta et al.^20^. (**H**) RT–qPCR analysis of microglial signature genes *CX3CR1*, *P2RY12*, *TMEM119*, *TREM2* and *MERTK* in HC and MS iMGLs. *n* = 8–12 wells, 4 HC cell lines and 6 MS cell lines, with 1–3 independent differentiations. The data are presented as single datapoints and medians and are normalized to the HC1 vehicle. Mann–Whitney U test. The grey values indicate the statistical significance of differences between the HC VEH and LPS groups, the green values between the MS VEH and LPS groups, and the black values between the HC and MS groups. Panel A was created in BioRender. Lotila, J. (2024) https://BioRender.com/k52w144.

Next, we compared the transcriptomes of our iMGLs with an scRNA-seq dataset from chronic active MS lesions published by from Absinta et al.^20^. Our analysis revealed that the iMGL most closely resembled the cluster annotated as “microglia inflamed in MS-iron” (MIMS-iron) located at the edge of chronic active lesions (Fig. 2G). Validating the expression of the microglial signature, the quantitative PCR (qPCR) analysis showed that iMGLs expressed *CX3CR1*, *P2RY12*, *TMEM119*, *TREM2*, and *MERTK* under basal conditions and exhibited downregulation of these markers in response to LPS treatment, except for *MERTK* (Fig. 2H). Notably, we observed a significant reduction in the expression of key homeostatic marker *P2RY12* in MS iMGLs compared with HC iMGLs under basal conditions (Fig. 2H). Collectively, our data indicate that the HC and MS iMGLs present the characteristic phenotypic traits of human microglia and resemble MS-associated microglia at the transcriptional level.

### Proinflammatory stimulation demonstrates the immune competence of the iMGLs

The transcription factor nuclear factor kappa-B (NF-κB) plays a central role in the pathogenesis of MS, mediating the activation of microglia^55^. We therefore wanted to test the immune competence and activation of this regulator of inflammation in iMGLs after stimulation with the proinflammatory factors IFN-γ (20 ng/ml), LPS (20 ng/ml) or their combination. Both HC and MS iMGLs rapidly responded to LPS, as evidenced by the nuclear translocation of NF-κB p65 after 15 min of exposure and a further increase after 45 min (Supplementary Fig. 4). The quantification of immunofluorescence staining after 45 min revealed NF-κB activation in a significant portion of iMGLs following LPS stimulation compared to vehicle-treated control cells (Fig. 3A and B). Compared with LPS alone, synergistic stimulation with LPS and IFN-γ led to similar levels of NF-κB activation in iMGLs. IFN-γ alone did not activate NF-κB in iMGLs. However, IFN-γ treatment altered the morphology, expression of microglial marker genes, and release of proinflammatory cytokines, confirming that iMGLs have the capacity to respond to several stimuli (Supplementary Fig. 5).

**Figure 3.**
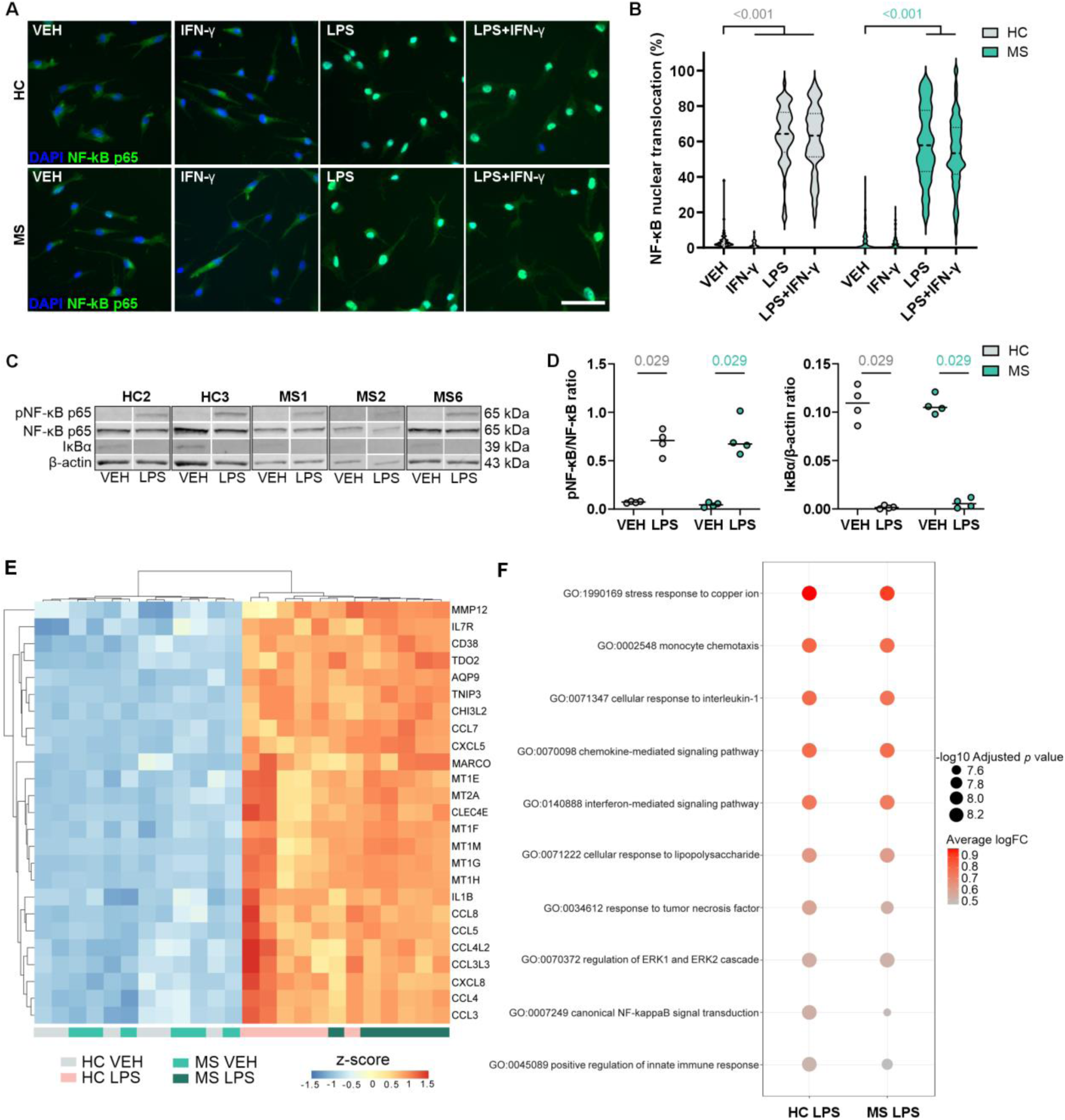
NF-κB activation and induction of inflammatory marker expression in LPS-treated iMGL. (**A**) Representative images of immunofluorescence staining for NF-κB p65 in vehicle-, IFN-γ-, LPS-or IFN-γ+LPS-stimulated HC and MS iMGLs. Scale bar = 50 µm. (**B**) Violin plot showing the quantification of the nuclear localization of NF-κB p65 from the immunofluorescence images. *n* = 54–118 images, 4 HC cell lines and 6 MS cell lines, with 2– 3 independent differentiations. Mann–Whitney U test. The Bonferroni post hoc correction was used for multiple comparisons. (**C**) Representative western blots showing the levels of phospho-NF-κB p65, NF-κB p65, IκBα and β-actin as a loading control from vehicle-and LPS-treated (45 min, 20 ng/ml) HC and MS iMGLs. (**D**) Quantification of the phospho-NF-κB p65 and NF-κB p6 ratios and IκBα levels normalized to β-actin. *n* = 4 wells, 2 HC cell lines and 3 MS cell lines, with 1–2 independent differentiations, each with 1–2 samples. The data are presented as single datapoints and medians. Mann–Whitney U test. See also Supplementary Fig. 6. (**E**) Heatmap of the hierarchical clustering analysis showing the top 20 shared DEGs with the highest logFC values in the LPS treatment group compared with the vehicle group. The colour corresponds to the z-score. (**F**) Dot plot depicting the selected shared (gseGO) terms from DEGs of LPS treatment. The size depicts the significance level, and the colour scale depicts the average logFC.

NF-κB activation was further investigated using western blotting, which allows an analysis of the phosphorylation of serine 536 on NF-κB p65 and the degradation of the inhibitory protein IκBa. As expected, quantification showed detection of phosphorylated NF-κB p65 after 45 min of LPS stimulation and, in contrast, IκBa levels were reduced, both of which are indicative of NF-κB pathway activation (Fig. 3C and D and Supplementary Fig. 6 and 7). However, no differences in the activation of NF-κB were observed between HC and MS iMGL.

Having shown the proinflammatory effect of LPS on NF-κB activation, we next sought to confirm the response at the transcriptome level after 24 h of treatment. The differential gene expression analysis revealed the greatest number of differentially expressed genes (DEGs; logFC ± 1 and adjusted *p* value < 0.05) between the LPS-and vehicle-treated groups, with 2,547 genes identified in HC iMGLs and 2,296 genes in MS iMGLs, confirming clear response to LPS. As anticipated, we observed the upregulation of numerous inflammatory genes in response to LPS treatment across different samples from both the HC and MS iMGL groups (Fig. 3E). Hierarchical clustering of the top shared DEGs revealed modules enriched in genes mediating inflammatory responses such as cytokines and chemokines and several metallothionine genes indicative of oxidative stress. Moreover, gene set enrichment analysis for Gene Ontology (gseGO) led to shared upregulation of processes related to the stress response to copper ion (*MT1G, MT1H, MT2A,* and *MT1M*), monocyte chemotaxis (*CCL7, CCL8, CCL5, CCL4,* and *CCL3*), chemokine-mediated signaling pathway (*CXCL8, CXCL5,* and *CCL7*) and canonical NF-κB signal transduction (*TNIP3, IL1B,* and *IL1A*) (Fig. 3F).

Taken together, these findings indicate that the generated iMGLs can elicit specific responses to different inflammatory stimuli and that the LPS-response confirms the reproducible induction of inflammatory marker expression in iMGLs.

### RNA sequencing reveals the upregulation of transcripts associated with immune activation in MS iMGL

Since microglia exhibit activated inflammatory phenotypes in rodent models of MS^21–23,56^ and in brain tissue samples from pwMS^20,21,27^, we next performed a comparative whole-transcriptome RNA-seq analysis to determine whether MS iMGLs display cell-intrinsic alterations compared with HC iMGLs. We analysed all six MS lines and all 4 HC lines under vehicle-and LPS-treated conditions. Remarkably, the DEG analysis confirmed the transcriptomic difference between MS and HC iMGLs in the basal state by identifying of 1,027 DE genes (using a cut-off of logFC > ± 1 and adjusted p value < 0.05; Fig. 4A). Furthermore, 154 genes were differentially expressed after LPS treatment between MS and HC iMGLs (Fig. 4A). Among these DEGs, 102 DEGs overlapped between the MS vehicle-treated group and the LPS-treated group (Fig. 4A).

**Figure 4.**
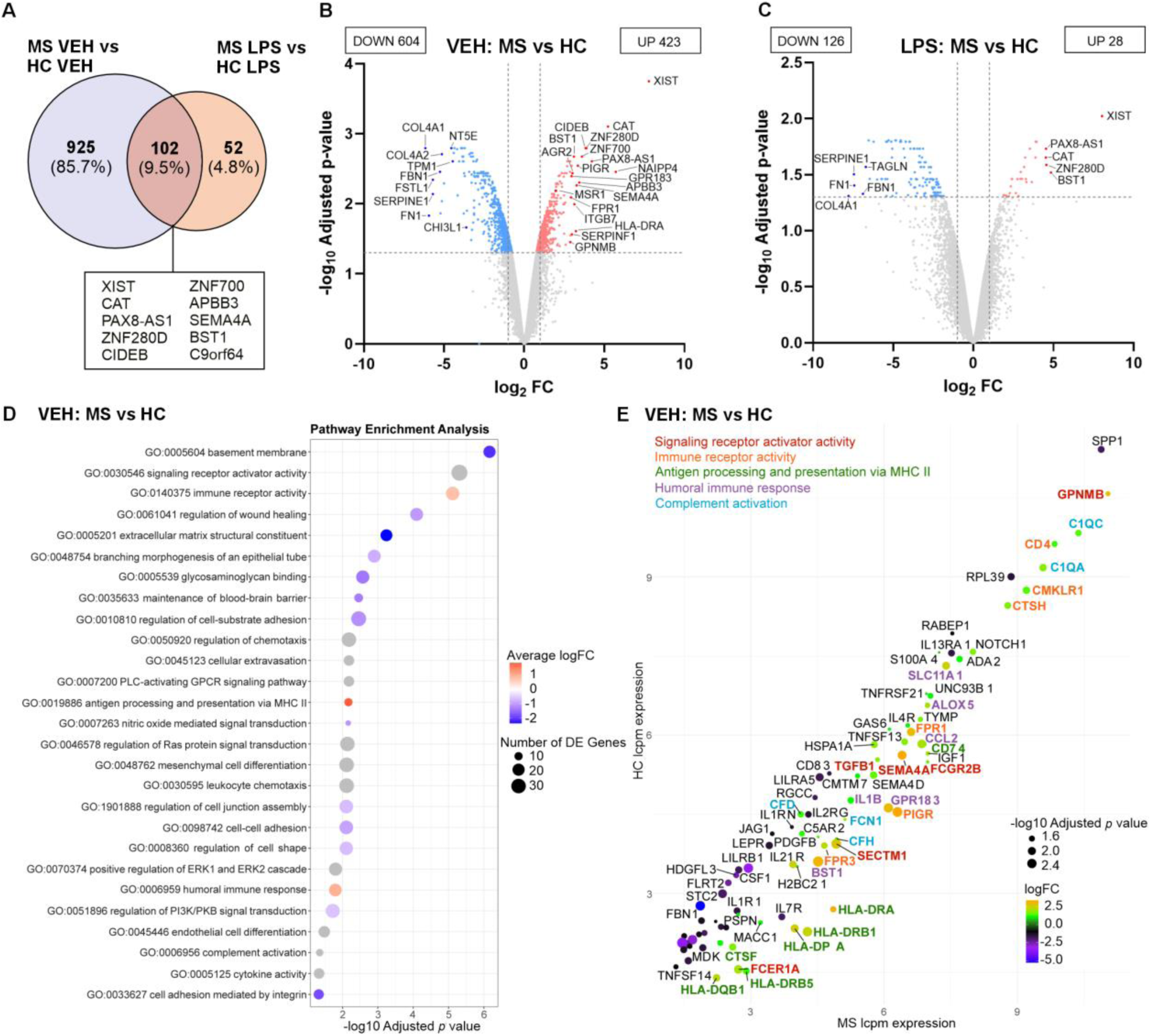
The RNA sequencing analysis revealed a shift in the immune activation state of MS iMGLs compared to HC iMGLs. (**A**) Venn diagram displaying the number of differentially expressed genes (DEGs) between MS and HC iMGLs stimulated with vehicle or LPS for 24 h, with adjusted *p* values < 0.05 and logFC values > ±1. Top overlapping upregulated DEGs between vehicle-and LPS-stimulated MS iMGLs. (**B**) Volcano plot showing DEGs identified between vehicle MS and HC iMGLs and (**C**) DEGs identified between LPS-stimulated MS and HC iMGLs. Volcano plots indicate upregulated (red) and downregulated (blue) DEGs with adjusted *p* values < 0.05 and logFC values > ±1. The top and selected significant DEGs based on logFC are mentioned. (**D**) Dot plot displaying selected top Gene Ontology (GO) terms for vehicle MS vs. HC iMGLs. The colour scale indicates the average logFC and the size indicates the number of DEGs in the GO set. (**E**) Dot plot of the top DEGs enriched in selected GO terms for the comparison of vehicle-treated MS and HC iMGLs. Each dot represents a gene, the size indicates the significance level, genes are colour-coded by logFC, and selected gene labels are colour coded by pathways. *n* = 1–2 samples per cell line, 4 HC cell lines and 6 MS cell lines, with 1–2 independent differentiations.

DE genes observed between MS and HC iMGLs under vehicle-and LPS-treated conditions are displayed in volcano plots (Fig. 4B and C). Among the top DEGs in the vehicle-treated condition, novel genes, including *ZNF280D, CIDEB, ZNF700, APBB3, PIGR, ITGB7, BST1, AGR2* and *SERPINF1*, were upregulated. Moreover, our analysis revealed that several noncoding transcripts, including pseudogenes and long noncoding RNAs (*XIST, NAIPP4,* and *PAX8-AS1*), were differentially expressed in MS iMGLs. Most of the top upregulated genes in the MS vs. HC iMGL comparison were shared between the vehicle-and LPS-treated groups (Fig. 4A-C). Other top DEGs in vehicle-treated MS iMGLs included genes previously described to be involved in MS pathology, such as *GPR183* (also known as *EBI2*) and *FPR1*^57–59^, or those directly linked to MS microglia, such as *CAT, SEMA4A, HLA-DRA, HLA-DPA1, GPNMB, SLC11A1, CD74, MSR1, ALOX5, HSPA1A*, *C1QA* and *FCGR2B*^20,21,25,27,60–63^.

We elucidated the biological functions involved in MS and the GO enrichment analysis of MS iMGLs highlighted significantly upregulated genes associated with immune responses in the basal state. These terms included “immune receptor activity” (*PIGR, FPR1, IL21R, FPR3, FCER1A, CTSH,* and *FCGR2B)*, “antigen processing and presentation of exogenous peptide antigen via MHC class II” (*HLA-DRA, HLA-DPA1, HLA-DRB1, HLA-DQB1, CD74,* and *HLA-DRB5*), “humoral immune response” (*BST1, GPR183, SLC11A1, ALOX5, CCL2,* and *IL1B*), and “complement activation” (*C1QA, FCN1,* and *C1QC*), among others (Fig. 4D). In addition, downregulated genes (including *COL4A1, FN1, SERPINE1, FSTL1, FBN1, COL4A2, TPM1,* and *CHI3L1*) were enriched for terms related to extracellular matrix organization, wound healing, cell adhesion and the regulation of cell shape (Fig. 4B and D). Similarly, LPS stimulation mainly downregulated these ECM-associated pathways in MS iMGLs compared with HC iMGLs.

Finally, we investigated five of the deregulated immune process-related pathways and their associated genes that influence the microglial phenotype. We compared the expression levels, fold change values and adjusted *p* values of DEGs between vehicle-treated MS iMGLs and HC iMGLs to identify the most significant genes (Fig. 4E). Interestingly, we distinguished several of the novel genes (*PIGR, BST1,* and *FPR3*) with immune functions and confirmed known MS-associated genes (*SEMA4A, HLA-DRA, HLA-DPA1, GPNMB, SLC11A1, CD74, ALOX5, HSPA1A*, *C1QA* and *FCGR2B*) that align with the active microglial state previously reported in various neurodegenerative diseases, including MS^20,21,25,27,60–63^.

In summary, these results reveal that MS iMGLs present intrinsic immune activation compared with HC iMGLs which is more evident in the basal state than after strong inflammatory stimulation with LPS. Furthermore, we identified the top DEGs and pathways related to antigen presentation, immune receptor activity and complement activation, indicating that increased microglial activation potentially plays a role in disease pathogenesis.

### MS iMGLs display altered cytokine secretion under basal and inflammatory conditions

As several immune-related pathways were upregulated in MS iMGLs these observations prompted us to investigate the release of inflammatory factors, a hallmark of microglial activation^15^. The release of various inflammatory cytokines and chemokines in the culture medium was analysed with multiplex assays following 24 h of vehicle or LPS stimulation. In the basal state, the secreted levels of TNF-α, IL-6, CXCL10, IL-1β, IL-10 and GM-CSF were negligible or lower than the detection range (<1 pg/ml), whereas the CXCL5 levels were low (<12 pg/ml), and the CXCL8 and CCL2 levels were considerable (median 200–400 pg/ml) (Fig. 5). Interestingly, MS iMGLs secreted more CCL2 in the basal state than HC iMGLs did (Fig. 5G), showing a pattern that was similar to the pattern of mRNA expression in the RNA-seq data (Fig. 4E).

**Figure 5.**
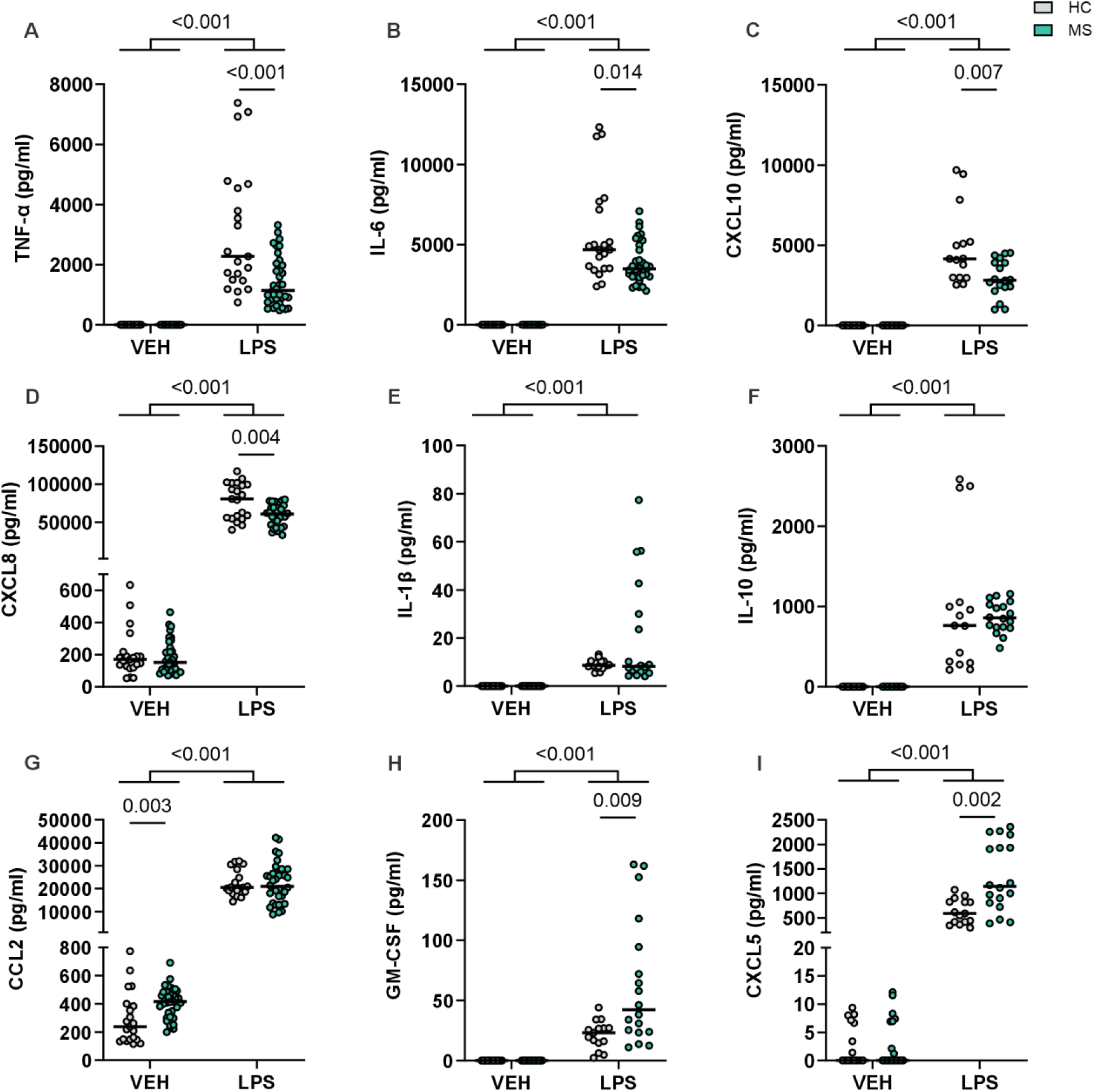
MS iMGLs exhibit altered cytokine release under basal and inflammatory conditions. Secretion of cytokines from HC and MS iMGLs stimulated with vehicle or LPS for 24 h. Multiplexed assay measuring (**A**) TNF-α, (**B**) IL-6, (**C**) CXCL10, (**D**) CXCL8, (**E**) IL-1β), (**F**) IL-10, (**G**) CCL2, (**H**) GM-CSF, and (**I**) CXCL5 levels in the media. *n* = 15–36 wells, 3 HC cell lines and 6 MS cell lines, with 1–2 independent differentiations per cell line, each with 3 samples. The data are presented as single datapoints and medians. Mann–Whitney U test.

LPS stimulation provoked a significant increase in the release of all measured cytokines and chemokines in both HC and MS iMGLs, indicating that LPS induced a switch from a surveying state to a proinflammatory microglial state. Compared with HC iMGLs, MS iMGLs released significantly higher levels of GM-CSF and CXCL5 (Fig. 5H and I). Concomitantly, MS iMGLs produced less TNF-α, IL-6, CXCL10 and CXCL8 as compared to HC iMGLs (Fig. 5A-D).

Thus, the generated iMGLs respond to the LPS stimulus, as expected, with increased cytokine and chemokine release. However, compared with HC iMGLs, MS iMGLs exhibit altered cytokine secretion both in the basal state and upon LPS stimulation.

### MS iMGLs show enhanced phagocytic function

Phagocytosis is a key function of microglia^16^; therefore, we explored whether MS iMGLs present functional changes in phagocytosis that are indicative of disease-promoting activity. Phagocytic function was evaluated using pHrodo zymosan A bioparticles and fluorescence imaging. We also tested the effects of proinflammatory pretreatment with IFN-γ, LPS or their combination on the phagocytosis of iMGLs. The spontaneous phagocytosis of the zymosan bioparticles was followed over 6 h (Supplementary Fig. 8A), and cytochalasin D treatment served as a negative control (Supplementary Fig. 8B and C). We observed phagocytosis in both HC and MS iMGLs, which was further modulated by inflammatory stimulation (Fig. 6A). Pretreatment of iMGLs with IFN-γ, and particularly with the combination of LPS+IFN-γ, greatly suppressed phagocytosis in all the studied HC and MS iMGLs (Fig. 6B). LPS pretreatment however, resulted in varying changes, with either a slight increase or decrease in phagocytosis in some of the HC and MS iMGL lines tested (Fig. 6B). The most notable finding was that phagocytic function was significantly increased in MS iMGLs compared to HC iMGLs, and while inflammatory stimulation modulated phagocytosis, MS iMGLs still exhibited significantly greater phagocytosis than HC iMGLs did (Fig. 6C). Thus, altered or excessive microglial phagocytosis may contribute to multiple sclerosis pathology.

**Figure 6.**
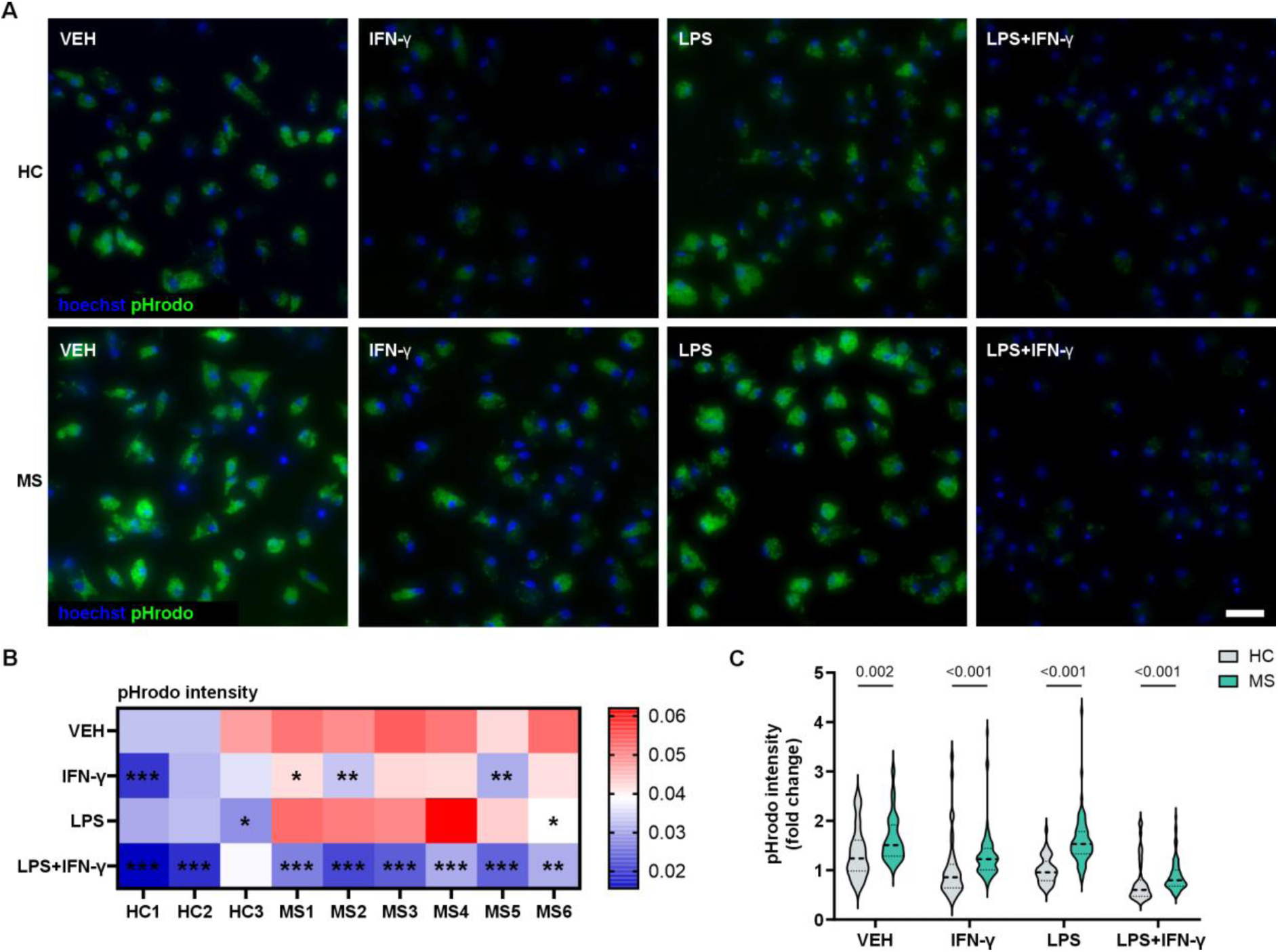
The phagocytic capacity of MS iMGLs is enhanced and modulated by inflammatory stimulation. (**A**) Representative fluorescence images of pHrodo zymosan bioparticles phagocytosed at 5 h after 24 h of vehicle, IFN-γ, LPS or IFN-γ+LPS stimulation in HC and MS iMGLs. Scale bar = 50 µm. (**B**) Heatmap indicating an increase (dark red) or decrease (dark blue) in the pHrodo fluorescence intensity normalized to the number of nuclei for each HC and MS iMGL cell line. *n* = 11–19 images per condition, with 2–3 differentiations per cell line. The heatmap presents the medians. Asterisks indicate statistically significant differences compared with the vehicle treatment within cell lines, as determined using the Mann–Whitney U test with Bonferroni’s correction \**p* < 0.05, \*\**p* < 0.01, and \*\*\**p* < 0.001. (**C**) Violin plot of the pHrodo intensity in HC and MS iMGL samples compared with the HC1 VEH sample. *n* = 21–79 images, 3 HC cell lines and MS 6 cell lines, with 2–3 differentiations per cell line. Mann–Whitney U test.

## Discussion

In this study, we investigated the intrinsic alterations of iPSC-based iMGLs derived from pwMS. We observed microglial activation using TSPO-PET imaging from progressed pwMS compared with the controls and successfully derived MS-specific iMGLs from these patients. Transcriptomic data revealed that MS iMGLs upregulated MS-associated genes and pathways related to immune regulation, antigen presentation and complement activation. This result was further confirmed by key functional assays showing an altered inflammatory secretome profile and increased phagocytic capacity.

In the context of MS, an increasing body of evidence indicates that the CNS compartmentalized inflammation driven by innate immune cells contributes to disease progression independently of relapse activity^7,10,14^. Both diffuse microglial activation in the NAWM and an increased chronic active lesion load, characterized by widespread microglial activation at the lesion edge, play significant roles in neurodegeneration^1,7^. In the present study, increased microglial activation was detected in pwMS using the TSPO-PET in the whole brain and the NAWM, which has been previously shown to predict later disease progression^14^. Therefore, we studied whether the intrinsic disease mechanisms of microglia can be recapitulated with iPSC-derived models. Here, iMGLs were successfully generated from these patients to introduce novel preclinical models of MS microglia. Although previous work by us and others using iPSC-derived glial and neuronal cells has shown that iPSCs provide a complementary platform to rodent models to study neuroinflammatory and neurodegenerative mechanisms *in vitro*^37,64^, to the best of our knowledge, the microglial phenotype has not been previously studied using iMGLs derived from pwMS.

First, our findings show that differentiated iMGLs closely correlate with the microglial signature observed in the previous studies of iPSC microglia, confirming the correct cell phenotype^38^. Interestingly, we were able to show that our differentiated iMGL resembled the transcriptional profile of microglia from MS lesions, particularly the cluster identified as MIMS-iron found at the chronic active lesion rim^20^. Additionally, our data revealed the downregulation of the homeostatic microglial gene *P2RY12* in MS iMGLs in the basal state. Consistent with our observation, earlier studies of postmortem tissues from patients with MS have indicated that microglia lose their homeostatic characteristic expression of *P2RY12*^20,24^. Our findings strongly support the use of the MS iMGL cell model as a powerful tool for revealing the cell-autonomous microglial alterations in pwMS.

Here, the transcriptional signature of MS iMGLs revealed the most biologically relevant changes occurring in the basal state without an external inflammatory stimulus, including the detection of novel immune-related transcripts such as *PIGR*, *BST1* and *FPR3*^65,66^. Interestingly, we also detected increased expression of the long noncoding RNA *XIST,* which has been recently described to contribute to inflammatory responses that potentially drive female-biased autoimmunity^67^. In addition, a previous snRNA-seq study of pwMS identified *XIST* among the top DEGs in a cluster annotated as macrophages at the chronic active lesion edge^20^, suggesting that *XIST* may be an important regulator of microglia and macrophage activation. Most importantly, MS iMGLs exhibited the upregulation of multiple genes previously reported to be expressed in MS microglia, such as *CAT, SEMA4A, HLA-DRA, HLA-DPA1, GPNMB, SLC11A1, CD74, MSR1, ALOX5, HSPA1A*, *C1QA* and *FCGR2B* ^20,21,25,27,60–63^. Among these, glycoprotein *GPNMB* is a known regulator of microglial immune responses, is linked to the phagocytic microglial phenotype and has been suggested as a marker for lesion-associated microglia^29,61,68^. Moreover, *HLA-DRA, HLA-DPA1, CD74, C1QA,* and *C1QC* levels were previously found to be elevated in MIMS-iron at the rim of chronic active lesions^20^. Therefore, these data demonstrate that MS iMGLs express several signature genes of reactive disease-associated microglial subtypes, as confirmed previously in both humans^20,21^ and in mice^21,69^.

Transcriptomic differences between MS and HC iMGLs in the basal state emphasized the role of microglia in immunoregulation, including the positive enrichment of pathways associated with immune receptor activity, antigen presentation and complement activation, among other processes, with known implications in MS^20^. The increased expression of MHC class II genes in our MS iMGLs is consistent with previous studies showing that in MS, activated microglia increase MHC molecules on their surface and can present antigens that activate T cells^20,70^. In addition, MS iMGLs upregulate complement genes compared with HC iMGLs, which is consistent with the involvement of the complement cascade in microglial activation and microglia-mediated synapse loss observed in MS^71^. Based on these findings, MS iMGLs present signs of cell-autonomous immune activation that may be key in CNS-confined MS pathology.

We examined these findings at the functional level and measured the release of inflammatory mediators from the culture media. The analysis confirmed increased secretion of CCL2 from MS iMGLs, which was in accordance with our RNA-seq data, suggesting that CCL2 contributes to microglial activation in the basal state. Prior reports of EAE and cuprizone mouse models of MS suggest that microglia are one of the primary sources of CCL2 in the brain^72,73^. Furthermore, a previous study using scRNA-seq of microglia from early active pwMS identified a cluster containing preactivated microglia with elevated CCL2 levels^21^. CCL2 has an important function in the recruitment of lymphocytes to the CNS and is recognized as a key regulator of glial function and microglial activation^72^. Moreover, in our study, LPS treatment activated the key inflammatory signalling cascade NF-κB in iMGLs, leading to changes in the secretion of several NF-κB targets^55^, including inflammatory cytokines and chemokines. Specifically, the secretion of GM-CSF and CXCL5 was increased following LPS challenge in MS iMGLs compared to HC iMGLs. Although GM-CSF plays an important role in microglial homeostasis, it also affects proliferation, antigen presentation, and phagocytosis^74^. CXCL5, the chemokine that attracts and activates neutrophils in response to autoimmune disorders^75^, has been shown to be expressed in immune-activated astrocytes^64^ and microglia^76^, and its release is attenuated by sphingosine receptor modulating MS drugs^76^. In contrast to a previous study of MS iPSC-derived astrocytes showing increased levels of a specific set of NF-κB targets following activation^34^, in our MS iMGLs, the levels of many of the tested cytokines, including TNF-α, IL-6, CXCL8 and CXCL10, were, in fact, decreased compared with those in HC iMGLs. A plausible explanation is that the reduced inflammatory response following LPS challenge represents an exhausted-like MG state, as our RNA-seq data, decreased *P2RY12* expression and increased CCL2 levels suggest that our MS iMGLs represent an already polarized immune state under basal conditions.

In MS, microglial phagocytosis plays a crucial role in the engulfment of myelin and clearance of cellular debris and contributes to synapse loss^25,61,71^. Here, we tested the phagocytic function of MS iMGLs and observed significantly enhanced phagocytosis both in the basal state and upon inflammatory challenge. Many of the upregulated genes that were detected in our MS iMGLs have also been linked to phagocytosis in transcriptional studies of postmortem tissues from pwMS^20,21,25^ suggesting that dysregulated phagocytosis contributes to MS pathology. Notably, a recent comprehensive analysis of microglial nodules from the NAWM revealed that activation by cytokines alongside phagocytosis of oxidized phospholipids likely contribute to a microglial phenotype that is predisposing to MS lesion formation^25^. The mechanism of microglial phagocytosis warrants further investigation, including how it impacts microglial activation, as this information will pave the way for the development of potential microglia-targeted treatment approaches.

Although our iPSC-derived MS microglial model allows us to elucidate cell type-specific disease mechanisms in a controlled setting, information on the effects of the complex three-dimensional cytoarchitecture and cell–cell interactions with other CNS or peripheral immune cells on microglial function is limited. Therefore, the development of more physiologically relevant cell models, such as the recently described glia-enriched organoid model^77^ incorporating MS iMGLs, would be necessary. Furthermore, even though we performed a thorough characterization of the four healthy control and six MS iMGL lines, adding more iPSC lines derived from patients presenting different disease types and levels of microglial activation would provide valuable insights into how the *in vitro* findings correlate with the *in vivo* observations.

In conclusion, these data demonstrate that microglia derived from pwMS exhibit cell-autonomous immune activation that may contribute to the neuroinflammatory milieu and pathological processes in MS. These findings indicate that microglial activation in the MS context can be studied with iPSC-derived iMGLs, and these cells represent a unique platform for the identification and testing of novel treatment targets.

## Data availability

The data that support the findings of this study are available within the article and its Supplementary material or are available from the corresponding author upon reasonable request. The raw RNA-seq data are not publicly available due to their containing information that could compromise the privacy of research participants. The RNA-seq data, including DEG lists and pathway analyses, will be shared upon publication.

## Acknowledgements

The authors acknowledge the services provided by Biocenter Finland (BF), the Tampere virus production facility, the Tampere Facility of iPS Cells, the Tampere Flow Cytometry Facility (TFCF), the Tampere Genomics Facility, the Tampere Imaging Facility (TIF) and the Tampere CellTech Laboratories (Tampere University). Moreover, the authors acknowledge Fimlab Laboratoriot Oy, Tampere, Finland, and the Institute for Molecular Medicine Finland FIMM Technology Centre, University of Helsinki, Finland, for their services. We thank Katriina Aalto-Setälä (Heart Group, Tampere University) for providing healthy control iPSC lines. We thank Eija Nieminen, Outi Melin, Hanna Pekkanen and Hanna Mäkelä for their technical assistance with iPSC production, cell maintenance and analyses. Furthermore, we thank Iikka Veijola, Alli Lintunen, Roosa Kattelus, Lassi Virtanen and Hanna Karvonen for their technical assistance with the analyses.

## References

1. Lassmann H. The contribution of neuropathology to multiple sclerosis research. European Journal of Neurology. 2022;29(9):2869–2877. doi:10.1111/ene.15360

2. Jakimovski D, Bittner S, Zivadinov R, et al. Multiple sclerosis. The Lancet. 2024;403(10422):183–202. doi:10.1016/S0140-6736(23)01473-3

3. Patsopoulos NA, Baranzini SE, Santaniello A. Multiple sclerosis genomic map implicates peripheral immune cells and microglia in susceptibility. Science (New York, NY). 2019;365(6460). doi:10.1126/science.aav7188

4. Harroud A, Stridh P, McCauley JL, et al. Locus for severity implicates CNS resilience in progression of multiple sclerosis. Nature. 2023;619(7969):323–331. doi:10.1038/s41586-023-06250-x

5. Lassmann H. Targets of therapy in progressive MS. Mult Scler. 2017;23(12):1593–1599. doi:10.1177/1352458517729455

6. Giovannoni G, Popescu V, Wuerfel J, et al. Smouldering multiple sclerosis: the “real MS.” Ther Adv Neurol Disord. 2022;15:17562864211066751. doi:10.1177/17562864211066751

7. Kuhlmann T, Moccia M, Coetzee T, et al. Multiple sclerosis progression: time for a new mechanism-driven framework. The Lancet Neurology. 2023;22(1):78–88. doi:10.1016/S1474-4422(22)00289-7

8. Kuhlmann T, Ludwin S, Prat A, Antel J, Brück W, Lassmann H. An updated histological classification system for multiple sclerosis lesions. Acta Neuropathol. 2017;133(1):13–24. doi:10.1007/s00401-016-1653-y

9. Frischer JM, Weigand SD, Guo Y, et al. Clinical and pathological insights into the dynamic nature of the white matter multiple sclerosis plaque. Ann Neurol. 2015;78(5):710–721. doi:10.1002/ana.24497

10. Absinta M, Sati P, Schindler M, et al. Persistent 7-tesla phase rim predicts poor outcome in new multiple sclerosis patient lesions. J Clin Invest. 2016;126(7):2597–2609. doi:10.1172/JCI86198

11. Dal-Bianco A, Grabner G, Kronnerwetter C, et al. Slow expansion of multiple sclerosis iron rim lesions: pathology and 7 T magnetic resonance imaging. Acta Neuropathol. 2017;133(1):25–42. doi:10.1007/s00401-016-1636-z

12. Airas L, Yong VW. Microglia in multiple sclerosis -pathogenesis and imaging. Curr Opin Neurol. 2022;35(3):299–306. doi:10.1097/WCO.0000000000001045

13. Oh U, Fujita M, Ikonomidou VN, et al. Translocator Protein PET Imaging for Glial Activation in Multiple Sclerosis. J Neuroimmune Pharmacol. 2011;6(3):354–361. doi:10.1007/s11481-010-9243-6

14. Sucksdorff M, Matilainen M, Tuisku J, et al. Brain TSPO-PET predicts later disease progression independent of relapses in multiple sclerosis. Brain: A Journal of Neurology. Published online 2020. doi:10.1093/brain/awaa275

15. Li Q, Barres BA. Microglia and macrophages in brain homeostasis and disease. Nature Reviews Immunology. 2018;18(4):225–242. doi:10.1038/nri.2017.125

16. Kettenmann H, Hanisch UK, Noda M, Verkhratsky A. Physiology of microglia. Physiol Rev. 2011;91(2):461–553. doi:10.1152/physrev.00011.2010

17. Paolicelli RC, Sierra A, Stevens B, et al. Microglia states and nomenclature: A field at its crossroads. Neuron. 2022;110(21):3458–3483. doi:10.1016/j.neuron.2022.10.020

18. Prinz M, Masuda T, Wheeler MA, Quintana FJ. Microglia and Central Nervous System– Associated Macrophages—From Origin to Disease Modulation. Annual Review of Immunology. 2021;39(Volume 39, 2021):251–277. doi:10.1146/annurev-immunol-093019-110159

19. Heppner FL, Greter M, Marino D, et al. Experimental autoimmune encephalomyelitis repressed by microglial paralysis. Nat Med. 2005;11(2):146–152. doi:10.1038/nm1177

20. Absinta M, Maric D, Gharagozloo M, et al. A lymphocyte–microglia–astrocyte axis in chronic active multiple sclerosis. Nature. 2021;597(7878):709–714. doi:10.1038/s41586-021-03892-7

21. Masuda T, Sankowski R, Staszewski O, et al. Spatial and temporal heterogeneity of mouse and human microglia at single-cell resolution. Nature. 2019;566(7744):388–392. doi:10.1038/s41586-019-0924-x

22. Nissen JC, Thompson KK, West BL, Tsirka SE. Csf1R inhibition attenuates experimental autoimmune encephalomyelitis and promotes recovery. Experimental Neurology. 2018;307:24–36. doi:10.1016/j.expneurol.2018.05.021

23. Ponomarev ED, Shriver LP, Maresz K, Dittel BN. Microglial cell activation and proliferation precedes the onset of CNS autoimmunity. Journal of Neuroscience Research. 2005;81(3):374–389. doi:10.1002/jnr.20488

24. Zrzavy T, Hametner S, Wimmer I, Butovsky O, Weiner HL, Lassmann H. Loss of “homeostatic” microglia and patterns of their activation in active multiple sclerosis. Brain: A Journal of Neurology. 2017;140(7):1900–1913. doi:10.1093/brain/awx113

25. van den Bosch AMR, van der Poel M, Fransen NL, et al. Profiling of microglia nodules in multiple sclerosis reveals propensity for lesion formation. Nat Commun. 2024;15(1):1667. doi:10.1038/s41467-024-46068-3

26. Singh S, Metz I, Amor S, van der Valk P, Stadelmann C, Brück W. Microglial nodules in early multiple sclerosis white matter are associated with degenerating axons. Acta Neuropathol. 2013;125(4):595–608. doi:10.1007/s00401-013-1082-0

27. Miedema A, Gerrits E, Brouwer N, et al. Brain macrophages acquire distinct transcriptomes in multiple sclerosis lesions and normal appearing white matter. Acta Neuropathologica Communications. 2022;10(1):1–18. doi:10.1186/S40478-021-01306-3/TABLES/1

28. Lerma-Martin C, Badia-i-Mompel P, Ramirez Flores RO, et al. Cell type mapping reveals tissue niches and interactions in subcortical multiple sclerosis lesions. Nat Neurosci. 2024;27(12):2354–2365. doi:10.1038/s41593-024-01796-z

29. Alsema AM, Wijering MHC, Miedema A, et al. Spatially resolved gene signatures of white matter lesion progression in multiple sclerosis. Nat Neurosci. Published online November 5, 2024:1–13. doi:10.1038/s41593-024-01765-6

30. Summers RA, Fagiani F, Rowitch DH, Absinta M, Reich DS. Novel human iPSC models of neuroinflammation in neurodegenerative disease and regenerative medicine. Trends in Immunology. Published online September 21, 2024. doi:10.1016/j.it.2024.08.004

31. Nicaise AM, Wagstaff LJ, Willis CM, et al. Cellular senescence in progenitor cells contributes to diminished remyelination potential in progressive multiple sclerosis. Proceedings of the National Academy of Sciences of the United States of America. 2019;116(18):9030–9039. doi:10.1073/pnas.1818348116

32. Lopez-Caraballo L, Martorell-Marugan J, Carmona-Sáez P, Gonzalez-Munoz E. iPS-Derived Early Oligodendrocyte Progenitor Cells from SPMS Patients Reveal Deficient In Vitro Cell Migration Stimulation. Cells. 2020;9(8):1803. doi:10.3390/cells9081803

33. Morales Pantoja IE, Smith MD, Rajbhandari L, et al. iPSCs from people with MS can differentiate into oligodendrocytes in a homeostatic but not an inflammatory milieu. PLoS One. 2020;15(6):e0233980. doi:10.1371/journal.pone.0233980

34. Ponath G, Lincoln MR, Levine-Ritterman M, et al. Enhanced astrocyte responses are driven by a genetic risk allele associated with multiple sclerosis. Nature Communications. 2018;9(1):1–9. doi:10.1038/s41467-018-07785-8

35. Ghirotto B, Oliveira DF, Cipelli M, et al. MS-Driven Metabolic Alterations Are Recapitulated in iPSC-Derived Astrocytes. Annals of Neurology. 2022;91(5):652–669. doi:10.1002/ana.26336

36. Nishihara H, Perriot S, Gastfriend BD, et al. Intrinsic blood–brain barrier dysfunction contributes to multiple sclerosis pathogenesis. Brain. 2022;145(12):4334–4348. doi:10.1093/brain/awac019

37. Konttinen H, Cabral-da-Silva MEC, Ohtonen S, et al. PSEN1ΔE9, APPswe, and APOE4 Confer Disparate Phenotypes in Human iPSC-Derived Microglia. Stem Cell Reports. 2019;13(4):669–683. doi:10.1016/j.stemcr.2019.08.004

38. Abud EM, Ramirez RN, Martinez ES, et al. iPSC-Derived Human Microglia-like Cells to Study Neurological Diseases. Neuron. 2017;94(2):278–293.e9. doi:10.1016/j.neuron.2017.03.042

39. Haenseler W, Sansom SN, Buchrieser J, et al. A Highly Efficient Human Pluripotent Stem Cell Microglia Model Displays a Neuronal-Co-culture-Specific Expression Profile and Inflammatory Response. Stem Cell Reports. 2017;8(6):1727–1742. doi:10.1016/j.stemcr.2017.05.017

40. Douvaras P, Sun B, Wang M, et al. Directed Differentiation of Human Pluripotent Stem Cells to Microglia. Stem Cell Reports. 2017;8(6):1516–1524. doi:10.1016/j.stemcr.2017.04.023

41. Kurtzke JF. Rating neurologic impairment in multiple sclerosis. Neurology. 1983;33(11):1444–1444. doi:10.1212/WNL.33.11.1444

42. Polman CH, Reingold SC, Banwell B, et al. Diagnostic criteria for multiple sclerosis: 2010 Revisions to the McDonald criteria. Annals of Neurology. 2011;69(2):292–302. doi:10.1002/ana.22366

43. Ojala M, Prajapati C, Pölönen RP, et al. Mutation-Specific Phenotypes in hiPSC-Derived Cardiomyocytes Carrying Either Myosin-Binding Protein C Or α-Tropomyosin Mutation for Hypertrophic Cardiomyopathy. Stem Cells International. 2016;2016:1684792.

44. Hongisto H, Ilmarinen T, Vattulainen M, Mikhailova A, Skottman H. Xeno-and feeder-free differentiation of human pluripotent stem cells to two distinct ocular epithelial cell types using simple modifications of one method. Stem Cell Res Ther. 2017;8(1):291. doi:10.1186/s13287-017-0738-4

45. Häkli M, Kreutzer J, Mäki AJ, et al. Electrophysiological Changes of Human-Induced Pluripotent Stem Cell-Derived Cardiomyocytes during Acute Hypoxia and Reoxygenation. Stem Cells Int. 2022;2022:9438281. doi:10.1155/2022/9438281

46. Lotila J, Hyvärinen T, Skottman H, Airas L, Narkilahti S, Hagman S. Establishment of a human induced pluripotent stem cell line (TAUi008-A) derived from a multiple sclerosis patient. Stem Cell Research. 2022;63:102865. doi:10.1016/j.scr.2022.102865

47. Ewels PA, Peltzer A, Fillinger S, et al. The nf-core framework for community-curated bioinformatics pipelines. Nat Biotechnol. 2020;38(3):276–278. doi:10.1038/s41587-020-0439-x

48. Robinson MD, Oshlack A. A scaling normalization method for differential expression analysis of RNA-seq data. Genome Biol. 2010;11(3):R25. doi:10.1186/gb-2010-11-3-r25

49. Giudice L, Mohamed A, Malm T. StellarPath: Hierarchical-vertical multi-omics classifier synergizes stable markers and interpretable similarity networks for patient profiling. PLOS Computational Biology. 2024;20(4):e1012022. doi:10.1371/journal.pcbi.1012022

50. Franzén O, Gan LM, Björkegren JLM. PanglaoDB: a web server for exploration of mouse and human single-cell RNA sequencing data. Database. 2019;2019:baz046. doi:10.1093/database/baz046

51. Ritchie ME, Phipson B, Wu D, et al. limma powers differential expression analyses for RNA-sequencing and microarray studies. Nucleic Acids Res. 2015;43(7):e47. doi:10.1093/nar/gkv007

52. Yu G, Wang LG, Han Y, He QY. clusterProfiler: an R Package for Comparing Biological Themes Among Gene Clusters. OMICS. 2012;16(5):284–287. doi:10.1089/omi.2011.0118

53. Liberzon A, Subramanian A, Pinchback R, Thorvaldsdóttir H, Tamayo P, Mesirov JP. Molecular signatures database (MSigDB) 3.0. Bioinformatics. 2011;27(12):1739–1740. doi:10.1093/bioinformatics/btr260

54. Rissanen E, Tuisku J, Rokka J, et al. In Vivo Detection of Diffuse Inflammation in Secondary Progressive Multiple Sclerosis Using PET Imaging and the Radioligand ^11^C-PK11195. J Nucl Med. 2014;55(6):939–944. doi:10.2967/jnumed.113.131698

55. Guo Q, Jin Y, Chen X, et al. NF-κB in biology and targeted therapy: new insights and translational implications. Sig Transduct Target Ther. 2024;9(1):1–37. doi:10.1038/s41392-024-01757-9

56. Vainchtein ID, Alsema AM, Dubbelaar ML, et al. Characterizing microglial gene expression in a model of secondary progressive multiple sclerosis. Glia. 2023;71(3):588–601. doi:10.1002/glia.24297

57. Hsiao CC, Sankowski R, Prinz M, Smolders J, Huitinga I, Hamann J. GPCRomics of Homeostatic and Disease-Associated Human Microglia. Front Immunol. 2021;12:674189. doi:10.3389/fimmu.2021.674189

58. Wanke F, Moos S, Croxford AL, et al. EBI2 Is Highly Expressed in Multiple Sclerosis Lesions and Promotes Early CNS Migration of Encephalitogenic CD4 T Cells. Cell Rep. 2017;18(5):1270–1284. doi:10.1016/j.celrep.2017.01.020

59. Bihler K, Kress E, Esser S, et al. Formyl Peptide Receptor 1-Mediated Glial Cell Activation in a Mouse Model of Cuprizone-Induced Demyelination. J Mol Neurosci. 2017;62(2):232–243. doi:10.1007/s12031-017-0924-y

60. van der Poel M, Ulas T, Mizee MR, et al. Transcriptional profiling of human microglia reveals grey-white matter heterogeneity and multiple sclerosis-associated changes. doi:10.1038/s41467-019-08976-7

61. Hendrickx DAE, van Scheppingen J, van der Poel M, et al. Gene Expression Profiling of Multiple Sclerosis Pathology Identifies Early Patterns of Demyelination Surrounding Chronic Active Lesions. Front Immunol. 2017;8:1810. doi:10.3389/fimmu.2017.01810

62. Leitner DF, Todorich B, Zhang X, Connor JR. Semaphorin4A Is Cytotoxic to Oligodendrocytes and Is Elevated in Microglia and Multiple Sclerosis. ASN Neuro. 2015;7(3):1759091415587502. doi:10.1177/1759091415587502

63. Gray E, Kemp K, Hares K, et al. Increased microglial catalase activity in multiple sclerosis grey matter. Brain Res. 2014;1559:55–64. doi:10.1016/j.brainres.2014.02.042

64. Hyvärinen T, Hagman S, Ristola M, et al. Co-stimulation with IL-1β and TNF-α induces an inflammatory reactive astrocyte phenotype with neurosupportive characteristics in a human pluripotent stem cell model system. Scientific Reports. 2019;9(1):16944. doi:10.1038/s41598-019-53414-9

65. Yokoyama S. Genetic polymorphisms of bone marrow stromal cell antigen-1 (BST-1/CD157): implications for immune/inflammatory dysfunction in neuropsychiatric disorders. Front Immunol. 2023;14. doi:10.3389/fimmu.2023.1197265

66. Zhu J, Li L, Ding J, Huang J, Shao A, Tang B. The Role of Formyl Peptide Receptors in Neurological Diseases via Regulating Inflammation. Front Cell Neurosci. 2021;15:753832. doi:10.3389/fncel.2021.753832

67. Dou DR, Zhao Y, Belk JA, et al. Xist ribonucleoproteins promote female sex-biased autoimmunity. Cell. 2024;187(3):733–749.e16. doi:10.1016/j.cell.2023.12.037

68. Ripoll VM, Irvine KM, Ravasi T, Sweet MJ, Hume DA. Gpnmb is induced in macrophages by IFN-gamma and lipopolysaccharide and acts as a feedback regulator of proinflammatory responses. J Immunol. 2007;178(10):6557–6566. doi:10.4049/jimmunol.178.10.6557

69. Jordão MJC, Sankowski R, Brendecke SM, et al. Single-cell profiling identifies myeloid cell subsets with distinct fates during neuroinflammation. Science. 2019;363(6425):eaat7554. doi:10.1126/science.aat7554

70. Lodygin D, Odoardi F, Schläger C, et al. A combination of fluorescent NFAT and H2B sensors uncovers dynamics of T cell activation in real time during CNS autoimmunity. Nat Med. 2013;19(6):784–790. doi:10.1038/nm.3182

71. Werneburg S, Jung J, Kunjamma RB, et al. Targeted Complement Inhibition at Synapses Prevents Microglial Synaptic Engulfment and Synapse Loss in Demyelinating Disease. Immunity. 2020;52(1):167–182.e7. doi:10.1016/j.immuni.2019.12.004

72. Errede M, Annese T, Petrosino V, et al. Microglia-derived CCL2 has a prime role in neocortex neuroinflammation. Fluids Barriers CNS. 2022;19(1):68. doi:10.1186/s12987-022-00365-5

73. Janssen K, Rickert M, Clarner T, Beyer C, Kipp M. Absence of CCL2 and CCL3 Ameliorates Central Nervous System Grey Matter But Not White Matter Demyelination in the Presence of an Intact Blood–Brain Barrier. Mol Neurobiol. 2016;53(3):1551–1564. doi:10.1007/s12035-015-9113-6

74. Stanley ER, Biundo F, Gökhan Ş, Chitu V. Differential regulation of microglial states by colony stimulating factors. Front Cell Neurosci. 2023;17. doi:10.3389/fncel.2023.1275935

75. Wang LY, Tu YF, Lin YC, Huang CC. CXCL5 signaling is a shared pathway of neuroinflammation and blood–brain barrier injury contributing to white matter injury in the immature brain. Journal of Neuroinflammation. 2016;13(1):6. doi:10.1186/s12974-015-0474-6

76. O’Sullivan SA, O’Sullivan C, Healy LM, Dev KK, Sheridan GK. Sphingosine 1-phosphate receptors regulate TLR4-induced CXCL5 release from astrocytes and microglia. Journal of Neurochemistry. 2018;144(6):736–747. doi:10.1111/jnc.14313

77. Fagiani F, Pedrini E, Taverna S, et al. A glia-enriched stem cell 3D model of the human brain mimics the glial-immune neurodegenerative phenotypes of multiple sclerosis. Cell Reports Medicine. Published online August 8, 2024:101680. doi:10.1016/j.xcrm.2024.101680

