## Supplementary material for "Microglia from patients with multiple sclerosis display a cell-autonomous immune activation state"

#### Supplementary Figures

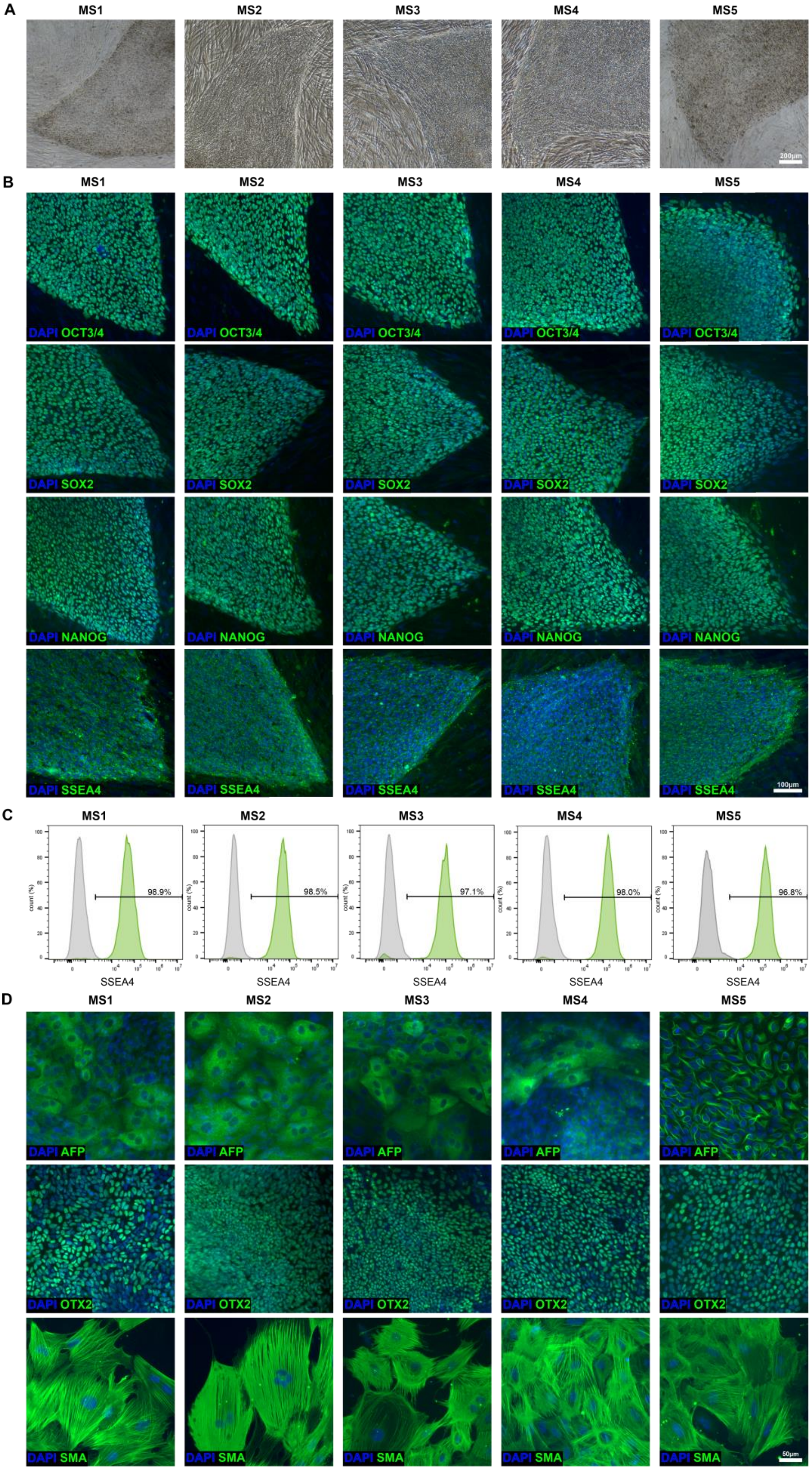
A
MS1
MS2
MS3
MS4
MS5
200μm
B
DAPI OCT3/4
DAPI SOX2
DAPI NANOG
DAPI SSEA4
100μm
C
98.9%
98.5%
97.1%
98.0%
96.8%
count (%)
SSEA4
D
DAPI AFP
DAPI OTX2
DAPI SMA
50μm

**Supplementary Figure 1. Characterization of the five MS patient-derived iPSC lines.** (**A**) Representative phase contrast images and (**B**) images of immunofluorescence staining for OCT3/4, SOX2, NANOG and SSEA4. (**C**) Flow cytometry analysis of SSEA4 expression. (**D**) Representative images of AFP, OTX2 and SMA immunofluorescence staining confirmed the capacity of iPSCs to differentiate into the three germ layers *in vitro* through embryoid body formation.

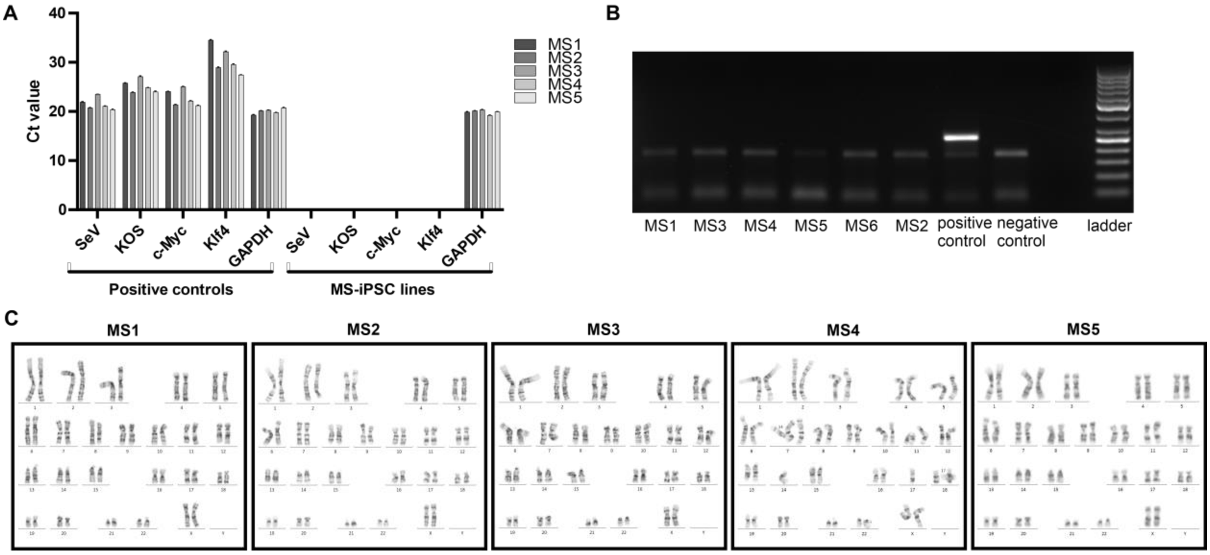


**Supplementary Figure 2. Characterization of MS patient-derived iPSC lines.** (**A**) RT–qPCR analysis of the removal of Sendai virus vectors and transgenes (SeV, KOS, c-Myc and Klf4) from MS iPSCs. Positive controls were collected after transduction (passage 0), and MS-iPSC samples were collected at passage 7–10. (**B**) Cells were negative for mycoplasma detected via RT–PCR. (**C**) The karyotype analysis confirmed the normal diploid 46,XX karyotype.

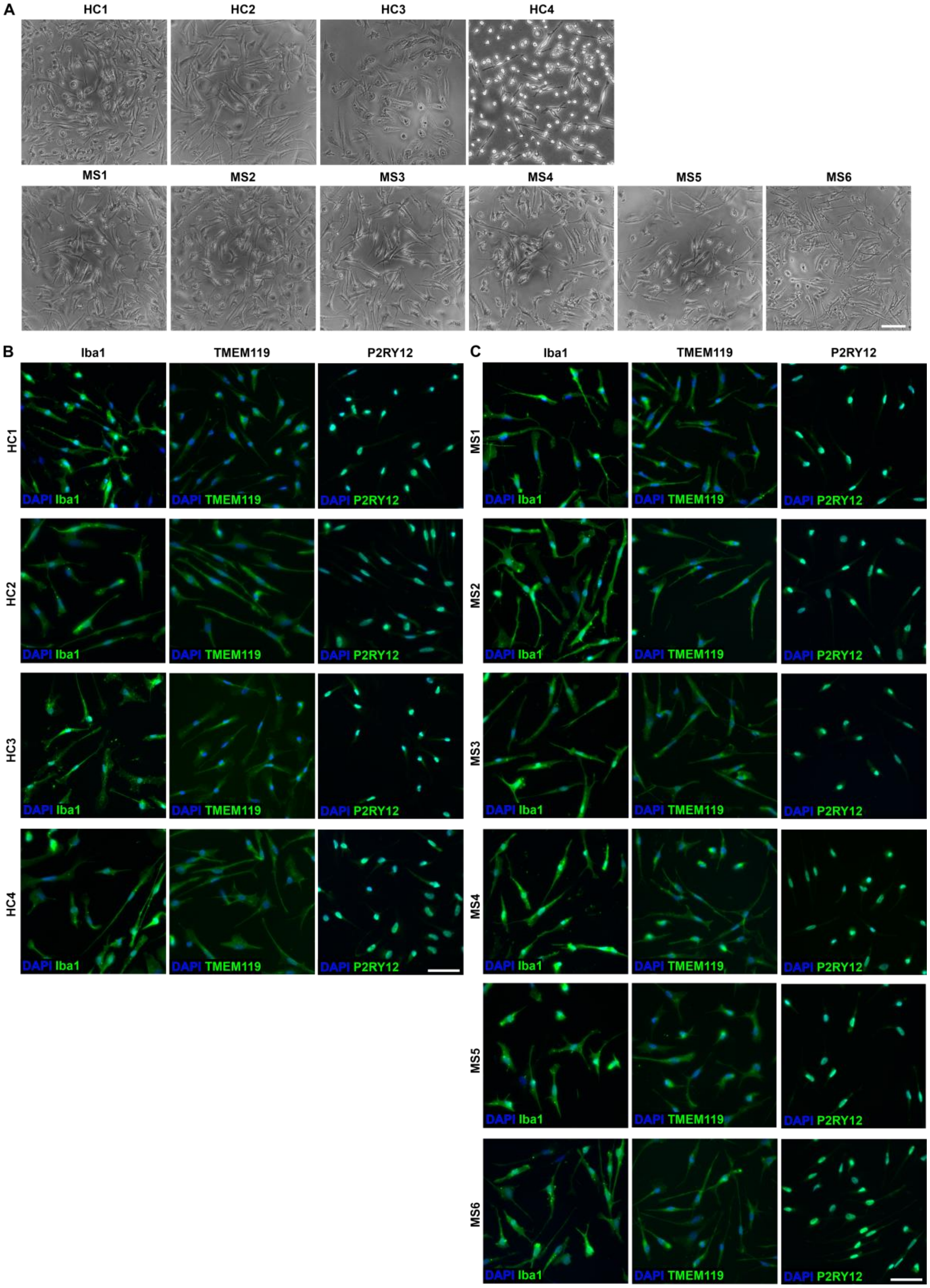


**Supplementary Figure 3. Characterization of HC and MS iPSC-derived iMGLs.** (**A**) Representative phase contrast images of HC and MS iMGLs. Scale bar = 100 µm. Representative images of immunofluorescence staining for Iba1, TMEM119 and P2RY12 in (**B**) HC and (**C**) MS iMGLs. Scale bar = 50 µm.

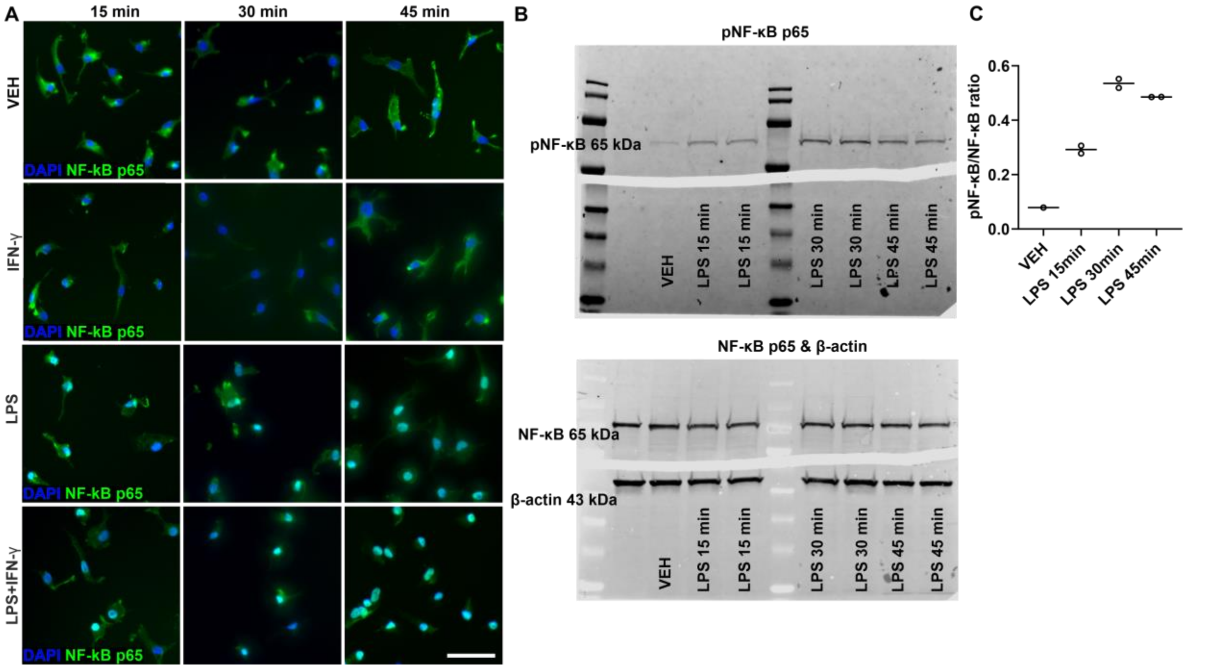


**Supplementary Figure 4. LPS stimulation activates the NF-κB signalling pathway in iMGLs.** (**A**) Representative images of immunofluorescence staining for NF-κB p65 at 15 min, 30 min and 30 min in unstimulated, IFNγ-, LPS- or IFNγ+LPS-stimulated HC and MS iMGLs. Scale bar = 50 µm. (**B**) Western blots showing the levels of phospho-NF-κB p65, NF-κB p65 and β-actin in unstimulated sample and samples treated with LPS for 15 min, 30 min or 45 min. Nine micrograms of protein was loaded into each lane. (**C**) Quantification of the phospho-NF-κB p65 and NF-κB p65 expression ratios. Protein levels were normalized to those of β-actin. *n* = 1–2 technical replicates (wells), with 1 differentiation. The data presented as single datapoints and medians.

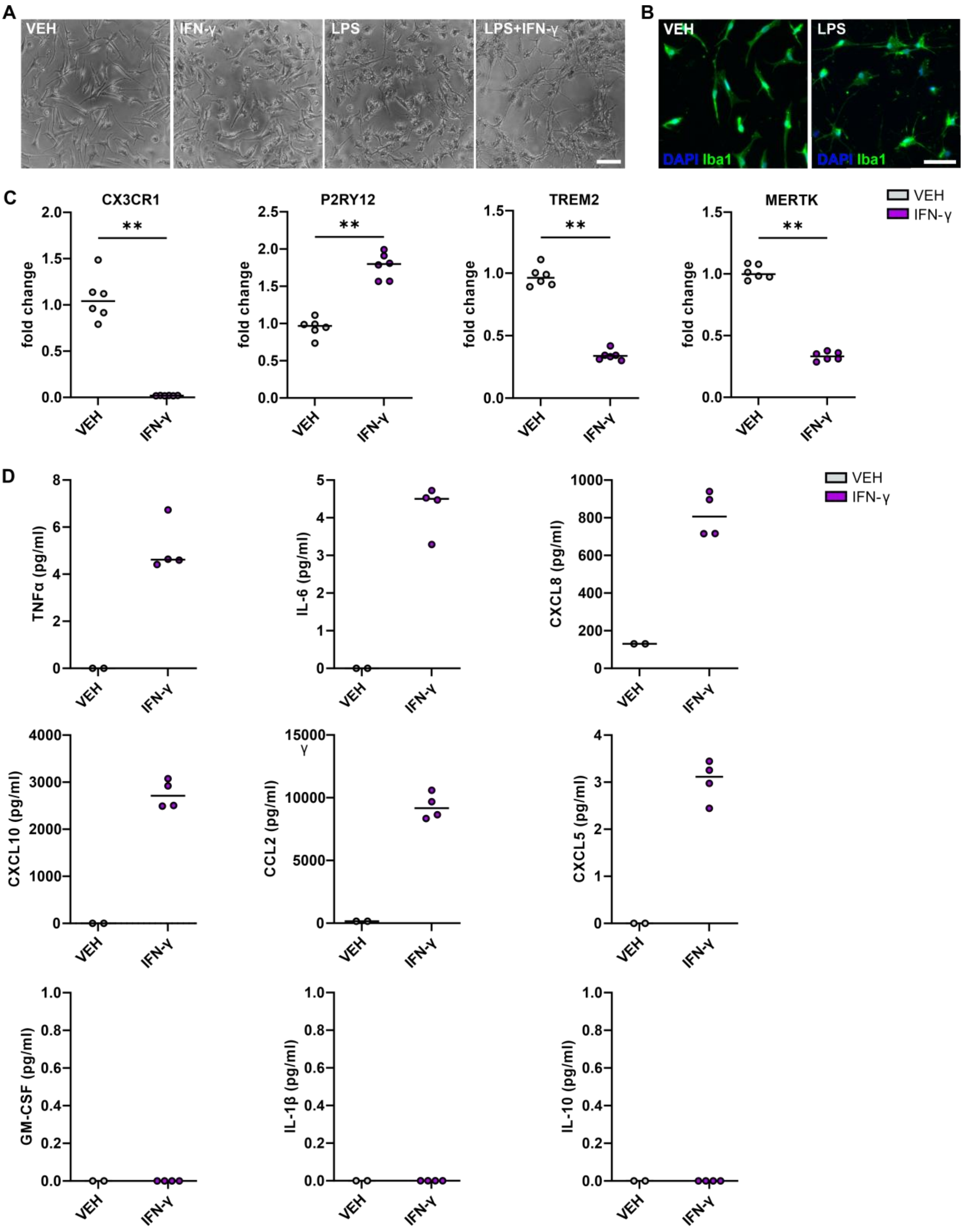


**Supplementary Figure 5. The effect of inflammatory stimulation on the iMGL phenotype.** (**A**) Representative phase contrast images of vehicle- and IFN-γ-, LPS- or IFN-γ+LPS-stimulated iMGLs after 24 h of treatment. Scale bar = 100 µm. (**B**) Representative images of Iba1 immunofluorescence staining in vehicle- and LPS-stimulated iMGLs after 24 h of treatment. Scale bar = 50 µm. (**C**) RT–qPCR analysis of the expression of the microglial signature genes *CX3CR1*, *P2RY12*, *TREM2* and *MERTK* in vehicle- and IFN-γ-stimulated iMGLs after 24 h of treatment ($n$ = 6 technical replicates, 1 HC cell line, with 1 differentiation). The data are presented as single data points and medians. (**D**) Analysis of cytokines secreted from vehicle- and IFN-γ-stimulated iMGLs after 24 h of treatment (1 HC cell line, with 1 differentiation). The data are presented as single data points and medians. Mann–Whitney U test, $^{**}p < 0.01$.

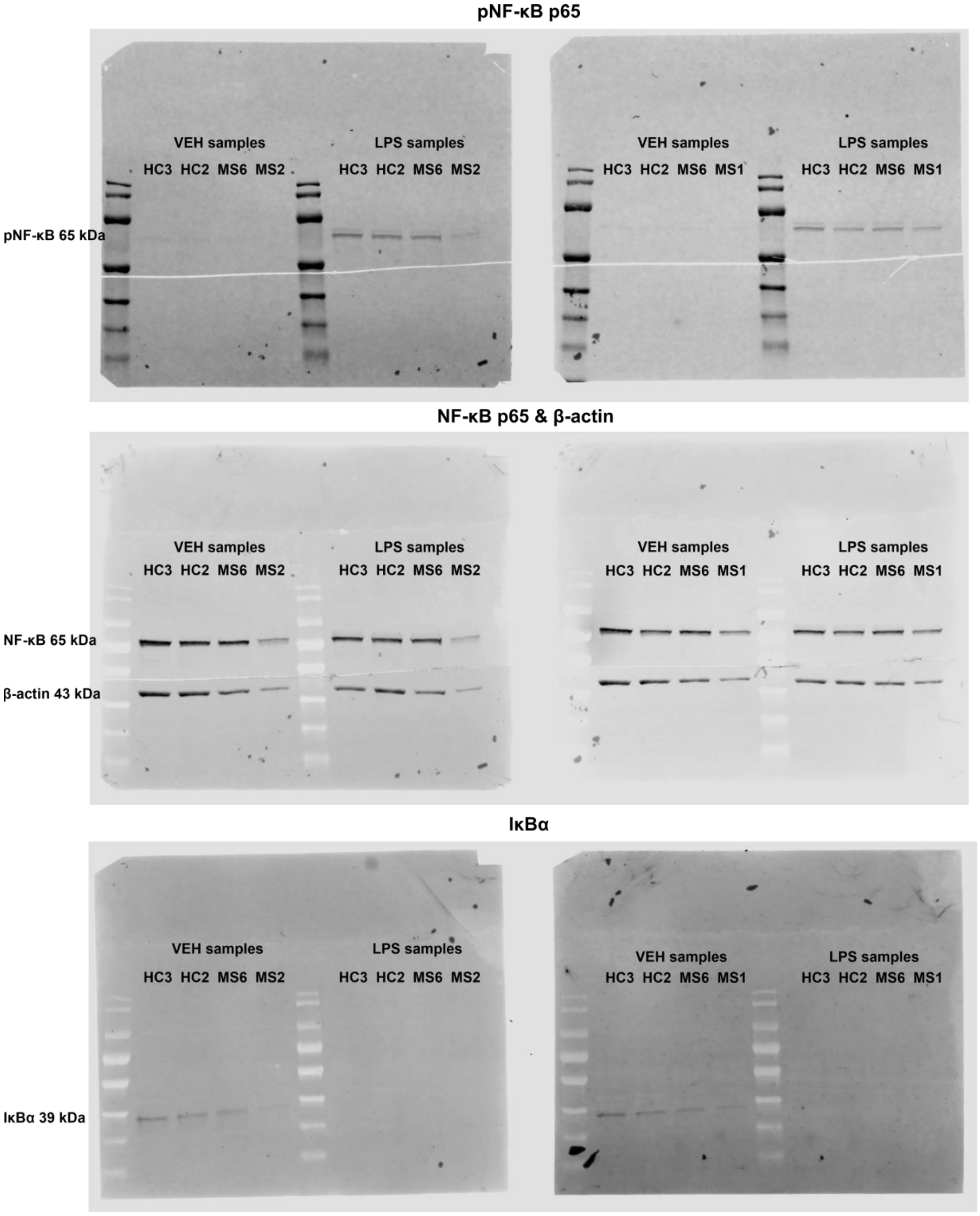

**Supplementary Figure 6. Western blots of NF-κB signalling proteins in HC and MS iMGL samples.** Western blots show the levels of phospho-NF-κB p65, NF-κB p65, β-actin and IκBα in vehicle- and LPS-treated (45 min, 20 ng/ml) HC (2 cell lines) and MS iMGLs (3 cell lines); 1-2 technical replicates (wells), 1-2 independent differentiations. Nine micrograms of protein was loaded per lane. See also Figure 3D.

A

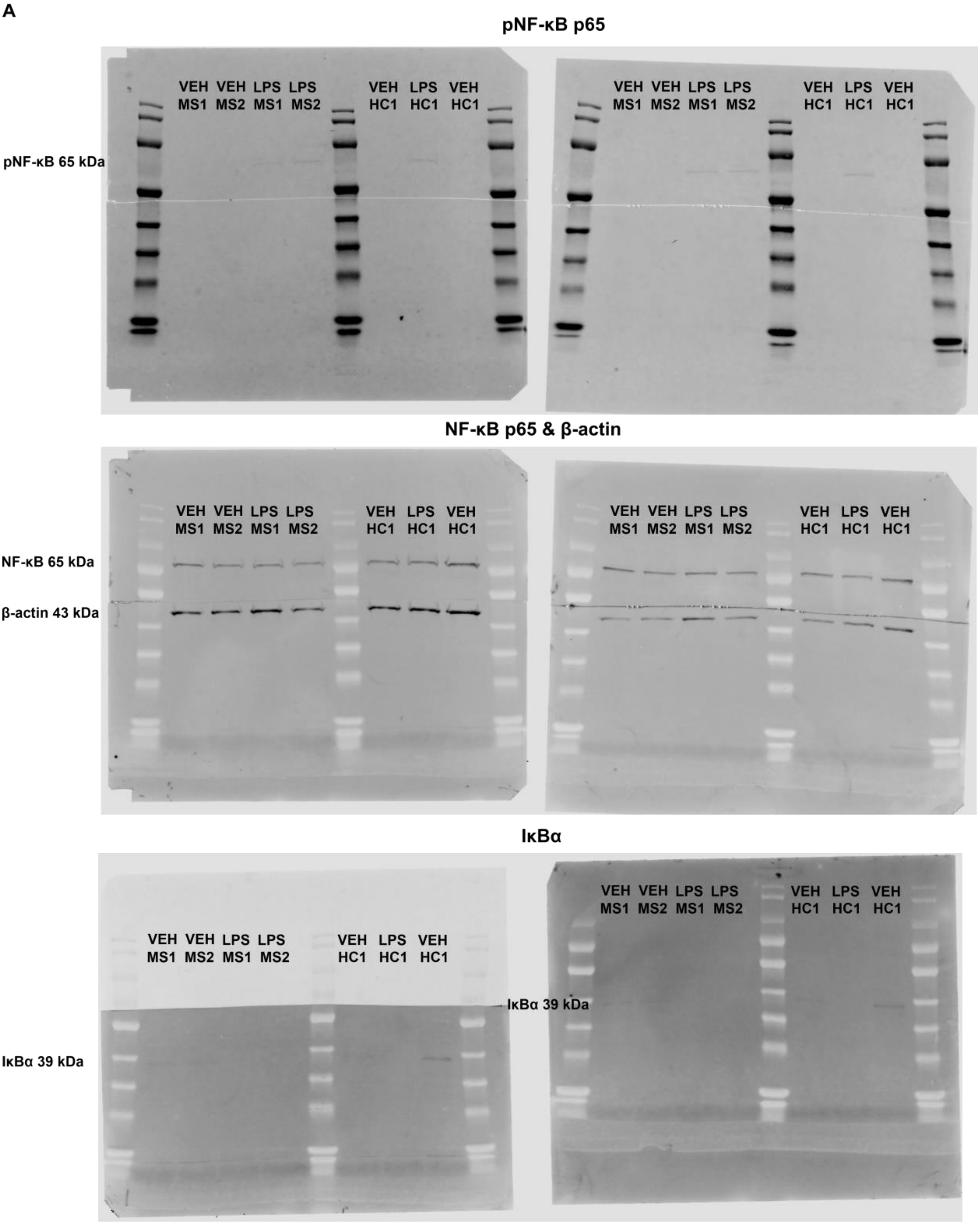
pNF-κB p65
pNF-κB 65 kDa
VEH MS1 VEH MS2 LPS MS1 LPS MS2 VEH HC1 LPS HC1 VEH HC1
NF-κB p65 & β-actin
NF-κB 65 kDa
β-actin 43 kDa
IκBα
IκBα 39 kDa

B

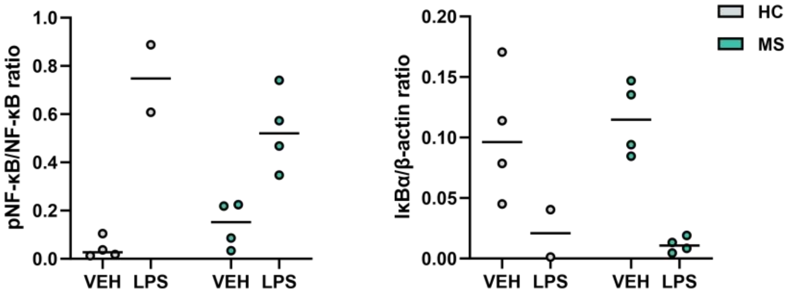
pNF-κB/NF-κB ratio
IκBα/β-actin ratio
VEH
LPS
HC
MS
0.0
0.2
0.4
0.6
0.8
1.0
0.00
0.05
0.10
0.15
0.20

**Supplementary Figure 7. Western blots of NF-κB signalling proteins in HC and MS iMGL samples.** (**A**) Western blots show the levels of phospho-NF-κB p65, NF-κB p65, β-actin and IκBα in vehicle- and LPS-treated (45 min, 20 ng/ml) HC (1 cell line, with 2 independent differentiations) and MS iMGLs (2 cell lines, with 1 differentiation). Five micrograms of protein was loaded per lane. (**B**) Quantification of the phospho-NFκB p65 and NFκB p65 expression ratios and IκBα protein levels. Protein levels were normalized to those of β-actin. *n* = 2–4 samples, with 2 independent differentiations. The data are presented as single datapoints and medians.

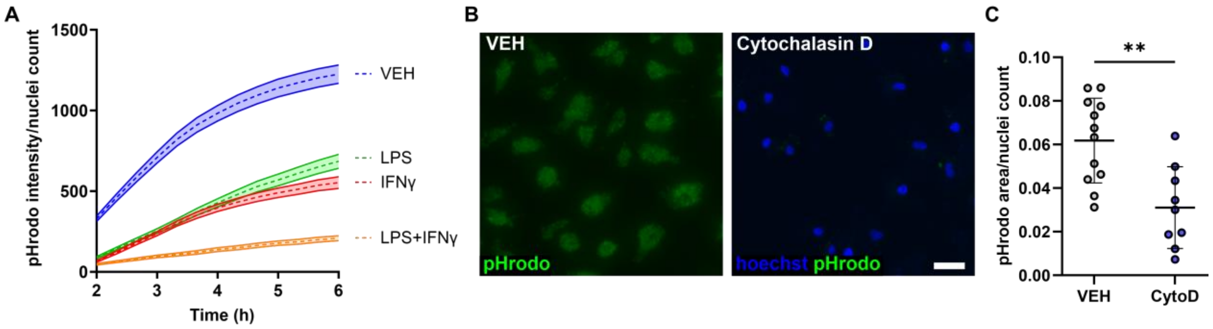


**Supplementary Figure 8. Analysis of iMGL phagocytosis.** (**A**) Representative time curve for the green fluorescence intensity of pHrodo zymosan phagocytosis in vehicle-, IFN-γ-, LPS- or IFN-γ+LPS-stimulated iMGLs after 24 h of treatment. *n* = 18 images per condition and time point; 6 wells/group, 1 differentiation. The data are presented as the means ± SEMs. (**B**) Representative fluorescence images of pHrodo zymosan bioparticles phagocytosed by iMGLs treated with the vehicle or the phagocytosis inhibitor cytochalasin D for 2 h. Scale bar = 50 µm. (**C**) Quantification of pHrodo zymosan phagocytosis in vehicle- and cytochalasin D-treated iMGLs. *n* = 9–12 images, 2–6 wells/group, with 2 independent differentiations. The data are presented as single data points and means ± SD. Unpaired t test, ***p* < 0.01.

### Supplementary Tables

**Supplementary Table 1.** Human induced pluripotent stem cell lines used in this study.

| Cell line | Alias | Sex | Karyotype | Health status | Reference |
|---|---|---|---|---|---|
| MS1 | MS1 | F | 46,XX | MS | this article |
| MS2 | MS2 | F | 46,XX | MS | this article |
| MS3 | MS3 | F | 46,XX | MS | this article |
| MS4 | MS4 | F | 46,XX | MS | this article |
| MS5 | MS5 | F | 46,XX | MS | this article |
| MS6 | MS6 | F | 46,XX | MS | Lotila et al. 2022[1] |
| UTA.04511.WTs | HC1 | M | 46,XY | Healthy | Ojala et al. 2016[2] |
| UTA.10902.EURCCs | HC2 | F | 46,XX | Healthy | Hongisto et al. 2017[3] |
| UTA.04602.WT | HC3 | F | 46,XX | Healthy | Ojala et al. 2016[2] |
| UTA.11311.EURCCs | HC4 | F | 46,XX | Healthy | Häkli et al. 2022[4] |

### Supplementary Materials

#### MRI and PET data acquisition and processing

Imaging was performed at the Turku University Hospital Neurocenter and Turku PET Centre. Conventional brain MRI using a 3T-MRI (Philips Ingenuity, Best, The Netherlands) was obtained with the following sequences: 3DT1, T2 and fluid-attenuated inversion recovery (FLAIR). Preliminary region of interest (ROI) masks for T1 lesions and FLAIR-based T2 lesions were created via the lesion segmentation toolbox (LST)[5] in SPM12 (Wellcome Trust Center for Neuroimaging, London, UK) and manually corrected after visual inspection to correspond to the T1 and T2 lesions. The whole brain, white matter (WM), grey matter (GM) and thalamic volumes were segmented after lesion filling using FreeSurfer (https://surfer.nmr.mgh.harvard.edu/). NAWM masks were obtained by subtracting the T2-lesion mask from the respective WM mask. Volumes of each ROI were evaluated as parenchymal fractions (PFs). Sixty-minute dynamic brain PET scans were performed using a brain-dedicated High-Resolution Research Tomograph scanner (HRRT, Siemens/Control Technology Incorporated, Knoxville, TN). The radiochemical synthesis of [$^{11}$C]PK11195, image reconstruction and post-processing were performed as previously described[6]. Each subject's PET image was co-registered with the respective T1 image using statistical parametric mapping (SPM8, Wellcome Trust Center for Neuroimaging). Specific binding of the radioligand was quantified as the distribution volume ratio (DVR). Time–activity curves corresponding to a reference region with no specific binding were obtained using Super-PK-software in MATLAB (MathWorks Inc.)[7], and Logan's reference tissue model[8] was applied for the DVR estimation.

#### Characterization of MS iPSC lines

MS iPSC lines were characterized as previously described[1]. Briefly, the expression of pluripotency markers (Oct3/4, Sox2, Nanog, and SSEA4) was analysed via immunofluorescence staining and flow cytometry. The trilineage differentiation capacity of iPSCs was determined *in vitro* by assessing embryoid body formation and performing immunofluorescence staining for endoderm (AFP), ectoderm (OTX2) and mesoderm (SMA) markers. The removal of Sendai virus vectors and transgenes (SeV, KOS, cMyc, and Klf4) was confirmed with RT–qPCR. The absence of mycoplasma was analysed using RT–PCR (Venor

GeM Classic Mycoplasma Detection Kit, Minerva Biolab). The karyotypes of the iPSC lines were analysed using the G-banding method at Fimlab Laboratoriot Ltd (Tampere, Finland). The genetic identities of the PBMCs and iPSCs were confirmed via a short tandem repeat (STR) analysis (GenePrint 24 system, Promega) at the Institute for Molecular Medicine Finland FIMM Technology Centre (University of Helsinki, Finland).

### Inflammatory treatments

The iMGLs were stimulated with lipopolysaccharide (LPS 0111:B4, 20 ng/ml, Sigma–Aldrich), interferon-γ (IFN-γ, 20 ng/ml, Peprotech), or a combination of LPS and IFN-γ (20 ng/ml for both) on Day 21 or 22. The iMGLs were stimulated for 45 min for NF-κB analyses and for 24 h for RNA-seq, RT–qPCR, secretion and phagocytosis assays. FBS starvation was performed 24 h prior phagocytosis and secretion assays simultaneously with inflammatory stimulation. Each differentiation included both healthy and MS iMGL lines, and the results consist of multiple independent differentiations as described in the figure legends.

### Immunocytochemistry

iMGLs were seeded on tissue-culture treated 96-well plates (PerkinElmer) at a density of 15,000 cells/well and fixed with 4% paraformaldehyde in phosphate-buffered saline (PBS) for 15 min. The cells were blocked with 10% normal donkey serum (NDS), 0.1% Triton X-100, and 1% bovine serum albumin (BSA) in PBS for 45 min at room temperature (RT). Primary antibodies were incubated with the cells in a solution containing 1% NDS, 0.1% Triton X-100, and 1% BSA in PBS overnight at 4 °C. The following primary antibodies were used: Iba1 (rabbit, 1:500; 019-19741; FujiFilm Wako), P2RY12 (rabbit, 1:125; HPA014518; Sigma-Aldrich), TMEM119 (rabbit, 1:100; ab185333; Abcam) and NF-κB p65 (rabbit, 1:400; D14E12; Cell Signaling Technology). The cells were incubated with an Alexa Fluor 488-conjugated donkey anti-rabbit secondary antibody (1:400; A21206; Thermo Fisher) diluted in 1% BSA in PBS for 1 h at RT. The stained cells were mounted with ProLong™ Gold Antifade Mountant with DAPI (Thermo Fisher Scientific) and imaged with an Olympus IX51 fluorescence microscope equipped with a Hamamatsu ORCA-Flash4.0 LT + sCMOS camera (type C11440-42U30). Both Iba1-positive iMGLs and NF-κB p65 nuclear translocation were quantified with CellProfiler (v4.2.1) and CellProfiler Analyst (v3.0.4)[9]. The NF-κB p65 analysis included three images per well and three wells per stimulation group. Images with low cell numbers (≤10 cells) were excluded from the analysis.

#### RT–qPCR

iMGLs were cultured on 6-well tissue culture-treated plates at a density of 400,000 cells/well. RNA was extracted from vehicle- and LPS-stimulated iMGLs after 24 h of treatment with a NucleoSpin RNA Kit (Macherey-Nagel) according to the manufacturer's protocol. The RNA was reverse transcribed to cDNA with a High Capacity cDNA Reverse Transcription Kit (Thermo Fisher Scientific). The mRNA expression levels of *CX3CR1* (Hs04187059_m1), *P2RY12* (Hs00224470_m1), *TMEM119* (Hs01938722_u1), *TREM2* (Hs00219132_m1) and *MERTK* (Hs01031979_m1) were determined with TaqMan assays utilizing an ABI QuantStudio 12K Flex Real-Time PCR System (Thermo Fisher Scientific). The data were analysed using the ΔΔCt method, with *GAPDH* (Hs99999905_ m1) serving as an endogenous control, and the data were normalized to those of thes HC1 vehicle sample.

#### Western blot analysis

iMGLs were cultured on 6-well tissue culture-treated plates at a density of 400,000 cells/well and treated with vehicle or LPS. The cells were washed with ice-cold PBS, and two wells were pooled with lysis buffer containing 50 mM Tris-HCl (pH 7.5), 10% glycerol, 150 mM NaCl, 1 mM EDTA, 50 mM NaF and 1% Triton X-100 (Sigma-Aldrich) on ice. Lysis buffer was supplemented with protease and phosphate inhibitor cocktails (both from Bimake). Lysed cells were centrifuged at 20 000 × g for 20 min at 4 °C. Protein concentrations were determined using Pierce™ 660 nm Protein Assay Reagent (Thermo Fisher Scientific) and 9 µg or 5 µg of protein was separated on 10% Mini-PROTEAN® TGX™ Precast Gels (456-1033, Bio-Rad) in Tris–Glycine–SDS running buffer. Proteins were transferred onto Trans-Blot® Turbo™ Mini PVDF Transfer membranes (#1704156, Bio-Rad) utilizing a Trans-Blot Turbo Transfer System (Bio-Rad). The membranes were blocked with 4% BSA in PBS for 1 h at RT, after which they were incubated with primary antibodies in 4% BSA in PBS overnight at 4 °C. The following primary antibodies were used: NF-κB p65 (1:1000, mouse, 6956; Cell Signaling Technology), phospho-NF-κB p65 (1:1000, rabbit, 3033; Cell Signaling Technology), IκBα (1:1000, mouse, 4814; Cell Signaling Technology) and β-actin (1:2000, mouse, sc-47778; Santa Cruz). Thereafter, the membranes were labelled with the secondary antibody in 4% BSA in PBS for 1 h at RT. The following secondary antibodies were used: IRDye® 800CW donkey anti-mouse IgG (1:20 000, 926-32212) and IRDye® 680RD donkey anti-rabbit IgG (1:20 000, 926-68073, both from LI-COR Biosciences). Proteins were detected with LI-COR Odyssey CLx imaging system, images were quantified with Image Studio software (both from LI-COR

Biosciences), and protein levels were normalized to those of β-actin. Normalized protein levels were used to quantify the ratio of phospho-NF-κB p65 to NF-κB p65.

#### Phagocytosis assay

iMGLs were cultured on 96-well tissue culture-treated plates (PerkinElmer) at a density of 15,000 cells/well and treated with vehicle, LPS, IFN-γ, or LPS+IFN-γ. After 24 h, the phagocytic capacity of iMGL was studied using pHrodo™ Green Zymosan Bioparticles™ Conjugate for Phagocytosis (P35365, Thermo Fischer Scientific). pHrodo bioparticles were added to the cells at a concentration 100 µg/ml in Opti-MEM (Thermo Fischer Scientific) supplemented with MCSF and IL-34 (both 10 ng/ml) and incubated for 6 h. The nuclei were stained with Hoechst 33342 at the end of the experiment (1:1000, Thermo Fischer Scientific). The cells were imaged (1 image/well, 6 wells/group) with an Olympus IX51 fluorescence microscope equipped with a Hamamatsu ORCA-Flash4.0 LT + sCMOS camera (type C11440-42U30). A Leica DMi8 inverted microscope was used for live-cell imaging every 20 min (5 %$CO_2$, 20% $O_2$, 37 °C; 3 images/well, 6 wells/group) to obtain a time curve after incubation with the pHrodo bioparticles for 2–6 h. The intensity and area of green fluorescence and the number of nuclei were analysed with CellProfiler[9] (v4.2.1). For the phagocytosis inhibition assay, iMGLs were pretreated with the actin polymerization inhibitor Cytochalasin D (10 µg/ml, Sigma–Aldrich) for 30 min prior to the addition of pHrodo bioparticles and imaged at 2 h.

#### Cytokine secretion

iMGLs were seeded on 96-well tissue culture-treated plates (PerkinElmer) at a density of 15,000 cells/well. The culture medium was pooled from two wells (n=3 samples/group in each experiment). The secretion of cytokines and chemokines (TNF-α, IL-6, IL-10, IL-1β, GM-CSF, CCL2, CXCL5, CXCL8, and CXCL10) in the vehicle-, LPS- or IFN-γ-treated iMGL cell culture medium was measured using a U-PLEX Custom Biomarker Group 1 (human) Assay (Meso Scale Diagnostics) according to the manufacturer’s instructions. The samples were run on a MESO QuickPlex SQ120 and analysed with DISCOVERY WORKBENCH® software (v4.0) (Meso Scale Diagnostics).

#### Statistical analysis

The normality of the data was analysed with the Shapiro–Wilk test. Normally distributed data were analysed with independent-sample t tests. Nonparametric Mann–Whitney U tests and Kruskal-Wallis tests were used for nonnormally distributed data. Bonferroni's or Dunn's post hoc test was used for multiple comparisons. A p value $< 0.05$ was considered to indicate statistical significance. Statistical analyses were performed with IMB SPSS Statistics software (version 29.0) and GraphPad Prism (version 10).

### References


1. Lotila J, Hyvärinen T, Skottman H, Airas L, Narkilahti S, Hagman S. Establishment of a human induced pluripotent stem cell line (TAUi008-A) derived from a multiple sclerosis patient. *Stem Cell Research*. 2022;63:102865. doi:10.1016/j.scr.2022.102865

2. Ojala M, Prajapati C, Pölönen RP, et al. Mutation-Specific Phenotypes in hiPSC-Derived Cardiomyocytes Carrying Either Myosin-Binding Protein C Or α-Tropomyosin Mutation for Hypertrophic Cardiomyopathy. *Stem Cells International*. 2016;2016:1684792.

3. Hongisto H, Ilmarinen T, Vattulainen M, Mikhailova A, Skottman H. Xeno- and feeder-free differentiation of human pluripotent stem cells to two distinct ocular epithelial cell types using simple modifications of one method. *Stem Cell Res Ther*. 2017;8(1):291. doi:10.1186/s13287-017-0738-4

4. Häkli M, Kreutzer J, Mäki AJ, et al. Electrophysiological Changes of Human-Induced Pluripotent Stem Cell-Derived Cardiomyocytes during Acute Hypoxia and Reoxygenation. *Stem Cells Int*. 2022;2022:9438281. doi:10.1155/2022/9438281

5. Schmidt P, Gaser C, Arsic M, et al. An automated tool for detection of FLAIR-hyperintense white-matter lesions in Multiple Sclerosis. *Neuroimage*. 2012;59(4):3774-3783. doi:10.1016/j.neuroimage.2011.11.032

6. Rissanen E, Tuisku J, Rokka J, et al. In Vivo Detection of Diffuse Inflammation in Secondary Progressive Multiple Sclerosis Using PET Imaging and the Radioligand $^{11}$C-PK11195. *J Nucl Med*. 2014;55(6):939-944. doi:10.2967/jnumed.113.131698

7. Turkheimer FE, Edison P, Pavese N, et al. Reference and target region modeling of [11C]-(R)-PK11195 brain studies. *J Nucl Med*. 2007;48(1):158-167.

8. Logan J, Fowler JS, Volkow ND, Wang GJ, Ding YS, Alexoff DL. Distribution volume ratios without blood sampling from graphical analysis of PET data. *J Cereb Blood Flow Metab*. 1996;16(5):834-840. doi:10.1097/00004647-199609000-00008

9. Stirling DR, Swain-Bowden MJ, Lucas AM, Carpenter AE, Cimini BA, Goodman A. CellProfiler 4: improvements in speed, utility and usability. *BMC Bioinformatics*. 2021;22(1):433. doi:10.1186/s12859-021-04344-9
